# Co-opted transposons help perpetuate conserved higher-order chromosomal structures

**DOI:** 10.1101/485342

**Authors:** Mayank NK Choudhary, Ryan Z Friedman, Julia T Wang, Hyo Sik Jang, Xiaoyu Zhuo, Ting Wang

**Author notes:** Corresponding author: Ting Wang, PhD, Department of Genetics, Washington University School of Medicine, 4515 McKinley Avenue, Campus Box 8510, St. Louis, MO 63110, USA.

## Abstract

Transposable elements (TEs) make up half of mammalian genomes and shape genome regulation by harboring binding sites for regulatory factors. These include architectural proteins—such as CTCF, RAD21 and SMC3—that are involved in tethering chromatin loops and marking domain boundaries. The 3D organization of the mammalian genome is intimately linked to its function and is remarkably conserved. However, the mechanisms by which these structural intricacies emerge and evolve have not been thoroughly probed. Here we show that TEs contribute extensively to both the formation of species-specific loops in humans and mice via deposition of novel anchoring motifs, as well as to the maintenance of conserved loops across both species via CTCF binding site turnover. The latter function demonstrates the ability of TEs to contribute to genome plasticity and reinforce conserved genome architecture as redundant loop anchors. Deleting such candidate TEs in human cells leads to a collapse of such conserved loop and domain structures. These TEs are also marked by reduced DNA methylation and bear mutational signatures of hypomethylation through evolutionary time. TEs have long been considered a source of genetic innovation; by examining their contribution to genome topology, we show that TEs can contribute to regulatory plasticity by inducing redundancy and potentiating genetic drift locally while conserving genome architecture globally, revealing a paradigm for defining regulatory conservation in the noncoding genome beyond classic sequence-level conservation.

**One-sentence summary:** Co-option of transposable elements maintains conserved 3D genome structures via CTCF binding site turnover in human and mouse.

## BACKGROUND

The 3D organization of various genomes has been mapped at high resolution using a variety of methods (*1-5*). While genome folding is largely conserved in mammals (*1,4*), the genetic forces shaping its emergence and evolution remain poorly understood. Two distinct yet mutually non-exclusive models (*6*) have recently gained much traction: that of phase separation (*7*) and of loop extrusion (*8,9*) by factors such as CTCF. In relation to the latter, TEs are known to contain and disseminate functional regulatory sequences (*10-13*) including that of CTCF. In contrast to relying on point mutations to evolve a functional CTCF binding site, TE transposition presents an attractive model for rapid regulatory sequence dissemination and regime building (*14-17*). Hence, we hypothesized that TEs have been a rich source of sequence for the assembly and tinkering of higher-order chromosomal structures. We studied the influence of all repetitive elements (REs) in establishing higher-order chromosomal structures and, more specifically, the role of TEs in the evolution of these higher-order chromosomal structures in humans and mice.

## RESULTS

We examined REs’ contribution to loop anchor CTCF sites using published genome-wide chromosomal conformation capture data from assays including ChIA-PET (*2*) and Hi-C in human (GM12878, HeLa, HMEC, IMR90, K562, NHEK) and mouse (ESCs, NSCs, CH12-LX) cell lines (*1*). We determined that 398 out of 3159 (12.6%) unique loop anchor CTCF sites were derived from REs in the mouse lymphoblastoid cell line. These RE-derived CTCF sites help establish 451 out of 2718 (16.6%) loops with discernible, unique CTCF loop anchors (Fig 1A, B). In the corresponding human lymphoblastoid cell line, REs contributed 935 out of 8324 (11.2%) unique loop anchor CTCF sites that help establish 1244 out of 8007 (15.6%) loops. Overall, REs contributed 9-15% of the anchor CTCF sites that result in 12-18% loops in humans and 12-23% of the anchor CTCF sites that result in 15-27% loops in mouse, across a variety of cell lines (Fig 1A, B).

**Figure 1:**
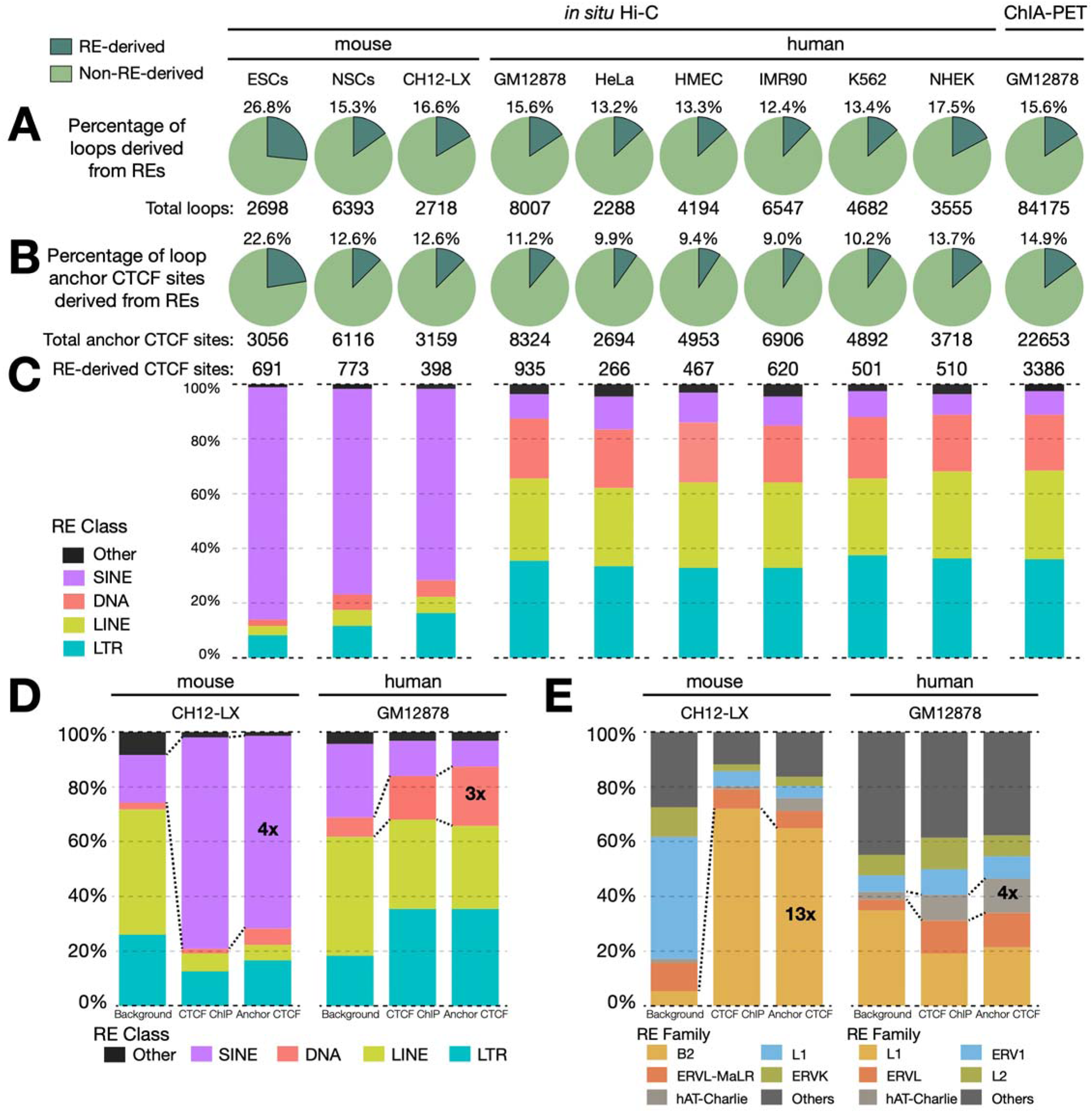
Contribution of repetitive elements (REs) to chromatin loops in humans and mouse. **(A)** Pie charts representing percentage of loops and (**B**) unique loop anchor CTCF sites derived from REs in a variety of human and mouse cell types. (**C**) Stacked bar plots showing the distribution of RE-derived anchor CTCF across major RE classes in the various human and mouse cell types. Stacked bar plots showcasing the distribution of RE-derived anchor CTCF vs. background and CTCF ChIP peaks across (**D**) major RE classes and (**E**) major RE families in matched blood lymphoblastoid cell line (mouse = CH12-LX; human = GM12878).

In both species, RE-derived loop anchor CTCF sites were largely derived from TEs (>95%) and their class of origin (SINE, LINE, LTR, DNA) showed a species-biased distribution (Fig 1C). Using the highest resolution *in-situ* HiC maps in matched lymphoblastoid cell types in mice (CH12-LX) and humans (GM12878), we compared the composition of the RE-derived loop anchor CTCF sites. While the mouse lineage was profoundly shaped by the SINEs (70%, 4x enrichment over background), the human lineage was overrepresented by retroviral LTR elements and DNA transposons (36% and 22%, 2x and 3x enriched over the background respectively) (Fig 1D). At the family level, the B2 SINEs in mice were 13-fold enriched over background and contributed 65% of TE-derived loop anchor CTCF sites. In humans, the hAT-Charlie family of DNA transposons contributed 13% of TE-derived loop anchor CTCF sites, a 4-fold enrichment over background (Fig 1E). These contributions are underestimates as we have yet to (i) uniquely identify all loop anchor CTCF sites (especially in repetitive regions), and (ii) annotate all repetitive elements, especially ancient TEs that have diverged far from their identity (*18*). Further, we looked at the cell-type specificity of these loop anchor CTCF sites in humans and see that 1334 out of 2017 (66%) RE-derived loop anchor CTCF sites were found in only one cell type (Supplementary Figure 1A). However, we did not find any specific TE family that enriches for cell-type specific loop anchor CTCF sites in the cell lines profiled (Supplementary Figure 1B).

To study the evolution of chromatin loops, we compared their conservation (Supplementary Methods) in matched human and mouse cell-types. Briefly, we used the liftOver tool (*19*) to compare loops across species and required exactly one reciprocal match (reciprocal best hit) to designate conserved loops. We found that 48% of all mouse loops (1596 out of 3331) had a loop call in the corresponding syntenic region in humans (Table S1.1). Our observation is in close agreement with prior studies (*1,4*) that show about half of all higher-order chromosomal structures to be conserved. We then sought to characterize the contribution of TEs to various classes of loops based on their orthology.

We compared the origin of loop anchor CTCF sites of orthologous loops in mouse and human. We found that out of 1596 orthologous loops, 142 (8.9%) in mouse and 108 (6.7%) in human had at least one TE-derived loop anchor CTCF site (Fig 2A). In addition to orthologous loops, TE-derived loop anchor CTCF sites also gave rise to 24% (409 out of 1735) and 15% (1136 out of 7852) non-orthologous (species-specific) loops in mouse and humans, respectively (Fig 2A), consistent with the appreciable role of TEs in genome innovation (*14-16,20,21*). Overall, the majority of TE-derived loop anchors in mouse were established by a handful of young TE subfamilies (B3, B2_Mm2, B3A, B2_Mm1t) that expanded in the rodent lineage (*22*) (Fig 2B). In contrast, multiple TE subfamilies of varying evolutionary ages contributed diffusely to CTCF loop anchors in humans (Fig 2C). Altogether, TEs in humans contributed to fewer orthologous loops and distributed over more TE subfamilies than in mouse.

**Figure 2:**
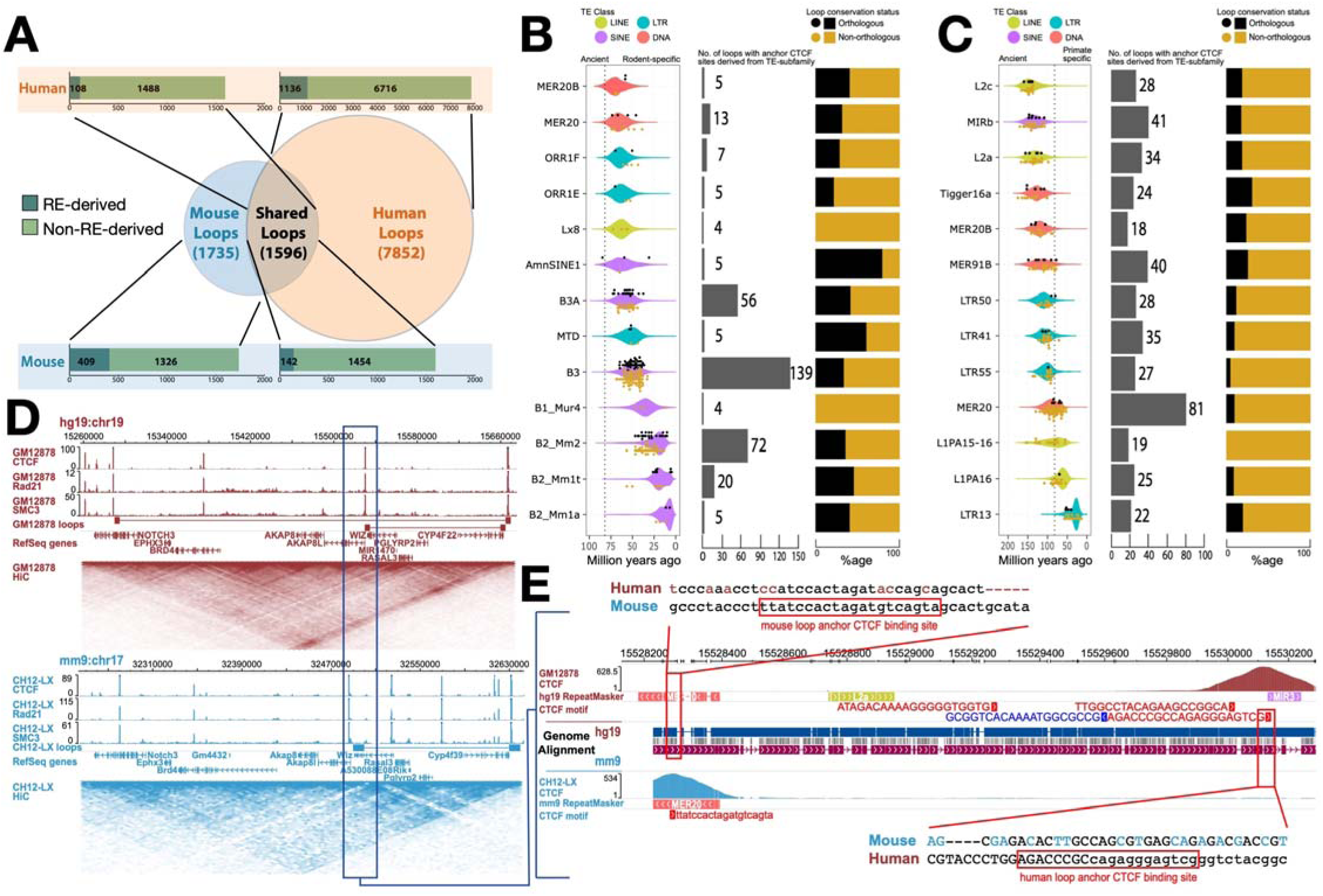
Contribution of TEs in the conservation landscape of human and mouse loops. **(A)** Venn diagram representing the various classes of chromatin loops based on their orthology and bar plots showing the contribution of REs to anchor CTCFs of each class of loops. (**B**) Age distribution and age of individual TEs that contribute loop anchor CTCF sites (black dots for orthologous loops; gold dots for non-orthologous loops) (left), total contribution to loop anchor CTCF sites (middle), distribution of orthologous and non-orthologous loops (right) derived from the top 13 TE subfamilies in mouse and (**C**) humans. Estimated primate/rodent divergence time (82 million years ago) is from (Meredith et al, 2011). (**D**) Contact maps representing a conserved chromatin loop in a syntenic region between human and mouse (**E**) A MER20 transposon insertion provides a redundant CTCF motif that helps in maintaining the conserved 3D structure via CTCF binding site turnover with remnants of the ancestral CTCF motif, well conserved in most non-rodent mammals (Supplementary Figure 2), still seen in the mouse genome.

Intriguingly, 123/142 (87%) TE-derived orthologous loops in mouse were discordant for TEs in humans (Table S1.2). In the sense: while the loops in humans were anchored at the putative ancestral CTCF binding sites, the syntenic ancestral CTCF motifs were largely degraded or deleted in mouse and the loops were now anchored at CTCF sites derived from nearby, co-opted TEs instead. One such example is an orthologous loop at the 5’ end of the Akap8l gene (Fig 2D) maintained in mouse by a MER20 element transposed ∼1.5kb upstream of the degraded ancestral motif which was well conserved in most non-rodent mammals (Supplementary Figure 2). The degradation of the ancestral CTCF motif derived from an ancient MIR3 element that is over 147 million-years-old (see *Methods*) incapacitates CTCF binding as evidenced by the CTCF-ChIP track (Fig 2E). In contrast, the younger MER20 element that inserted ∼90 million-years ago harbored strong CTCF binding, providing an anchor site to maintain the conserved loop in mouse. Similarly, we find that 89/108 (82%) TE-derived orthologous loops in human GM12878 cells were discordant for TEs in mouse (Table S1.3). We hypothesized that TEs provide redundant CTCF sites and mediated binding site turnover for CTCF contributing to conserved genome folding events between human and mouse.

Moreover, the 123 turned-over loops in mouse represent 127 turnover events (4 loops had both loop anchors turned-over) mediated by 124 unique loop anchors (3 turned-over loop anchors tethered 2 loops each). Out of the 124 unique loop anchors, 61 events represent turnover of the left loop anchor and 63 events represent turnover of the right loop. In terms of CTCF motif orientation—for the 61 left loop anchor turnover events, 53 were positive and 8 were negative; and for the 63 right loop anchor turnover events, 45 were negative and 18 were positive (Chi-square test, p-value=5.3×10^−11^). Similarly, in humans the 89 turned-over loops represent 93 turnover events (4 loops had both loop anchors turned-over) were mediated by 84 unique loop anchors (1 turned-over loop anchor tethered 3 loops, and 7 loop anchors tethered 2 loops each). Out of the 84 unique loop anchors, 43 events represent turnover of the left loop anchor (43 positive orientation CTCF motif and 0 negative orientation CTCF motif), and 41 events represent turnover of the right loop (40 positive orientation CTCF motif and 1 negative orientation CTCF motif) (Chi-square test, p-value=3.6×10^−19^). These results further lend credence to the loop extrusion model (*8*) and suggest that TE exaptation is more likely when the orientation of the inserted TE (and the underlying CTCF motif provided) is compatible with the local loop structure.

| **mm9 CH12-LX (n=124)** | Left Loop Anchor | Right Loop Anchor |
| --- | --- | --- |
| +ve CTCF motif | 53 | 18 |
| -ve CTCF motif | 8 | 45 |

| **hg19 GM12878 (n=84)** | Left Loop Anchor | Right Loop Anchor |
| --- | --- | --- |
| +ve CTCF motif | 43 | 1 |
| -ve CTCF motif | 0 | 40 |

Since the mouse genome is replete with repeat-derived CTCF sites (*22*) that could interfere with the targeted study of specific TE candidates, we decided to validate these hypotheses in human cell lines.

Here we examine two candidate TEs that maintain conserved higher-order chromosomal structures in humans: one belonging to the L1M3f subfamily of LINEs, and the other belonging to the LTR41 subfamily of endogenous-retrovirus-derived long terminal repeat (LTR). The former TE replaces the function of a lost ancestral CTCF site (Supplementary Figure 3), while the latter is functionally redundant for an ancestral CTCF site still present in humans (Supplementary Figure 4). These two TEs were specifically chosen as they could be unambiguously attributed to the genome folding function (no other CTCF/Cohesin binding site in the vicinity). Using CRISPR-Cas9, we obtained clones of GM12878 cells bearing homozygous deletions of the L1M3f and LTR41 elements, respectively (Supplementary Figure 5, Table S2.4). We then performed HYbrid-Capture on the *in situ* Hi-C library (Hi-C^2^) to examine the effect of the TE deletion on the local 3D structure (*8*) (Table S2.1, S2.2, S2.3).

The L1M3f-derived CTCF site was positioned at a conserved domain border and anchored three chromatin loops (Supplementary Figure 3). Upon deletion of this L1M3f, the conserved local chromosomal structure collapsed as evidenced by (i) the loss of focal enrichment in the homozygous TE knockout (KO) contact map in comparison to the wild-type (WT) contact map, and (ii) the fusion of two neighboring domains (Hi-C^2^ results: Fig 3A, Hi-C results: Supplementary Figure 6). The Virtual 4C plot anchored at the region surrounding the L1M3f element showed three distinct peaks (corresponding to the three loops in the WT cell line), which were lost in the KO (ΔL1M3f) cell line. We also found that cross-domain interactions significantly increased from 8% in WT to 19% in KO cell lines (∼2.4x, Welch’s t-test p-value<1.5×10^−16^, Table S2.5) across the L1M3f-established domain boundary, a change specific to the targeted domain and not seen in a control domain from a nearby region (Fig 3C). Thus, the L1M3f element is necessary for maintaining the conserved loops and domain boundary in humans. It represents a novel class of binding site turnover (*23-26*) for CTCF leading to conservation in terms of function via establishment of long-range interactions and potentially the underlying gene regulation, but not in primary local sequence.

**Figure 3:**
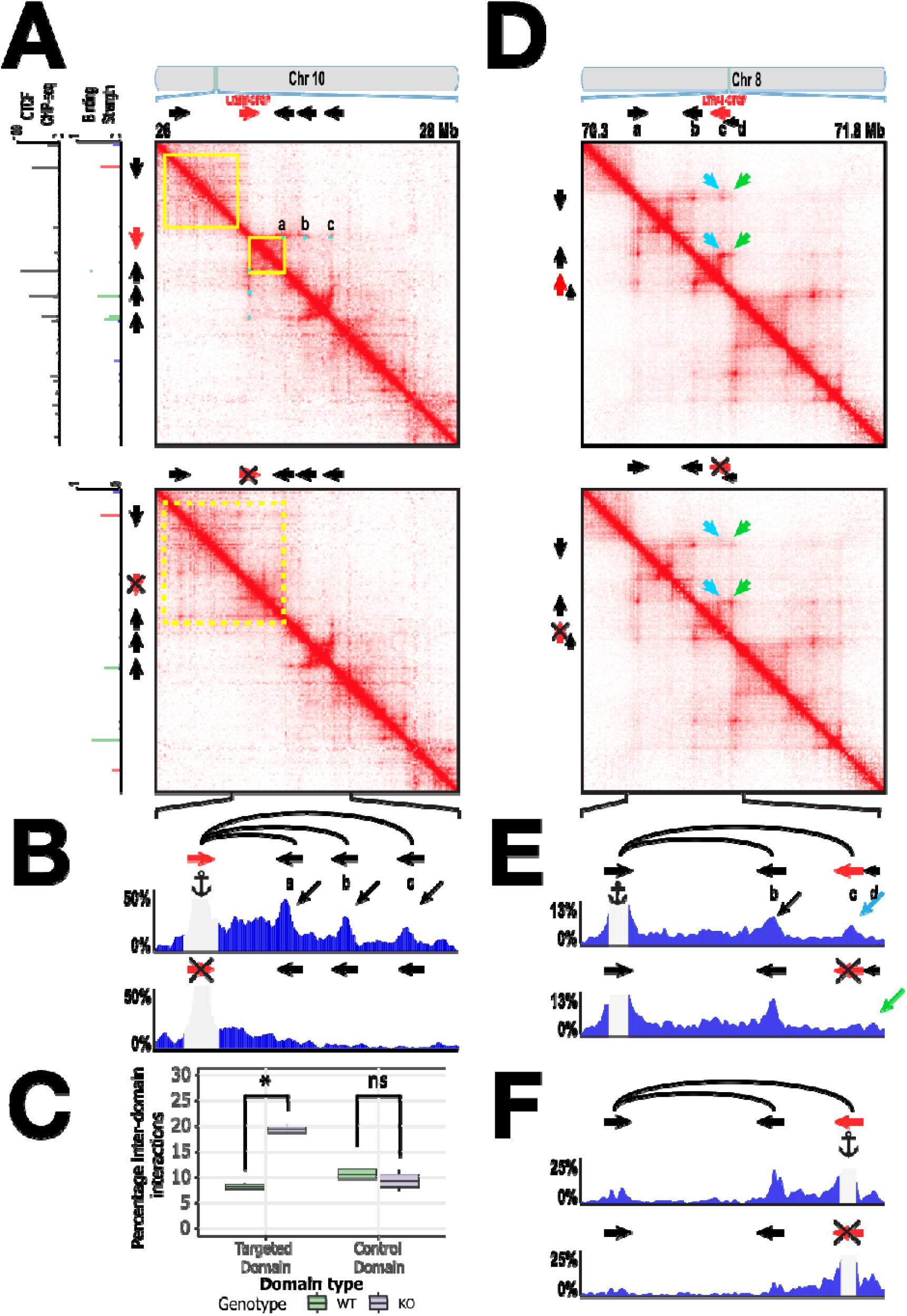
TEs are necessary for maintaining conserved higher-order chromosomal structures in humans. **(A)** Results of a CRISPR/Cas9-based deletion of an L1M3f element at chr10:26–28 Mb in GM187278 cells. Mega-contact maps (details in Methods) generated using Hi-C^2^ technology for the (top) WT locus and (bottom) KO (ΔL1M3f) locus. (**B**) Virtual 4C plot displaying total percent interactions emanating from an anchor on a 5kb-window containing the L1M3f element. (**C**) Boxplot measuring the percent inter-domain interactions (Table S2.5) across the targeted domain and a control domain (boundaries unaffected by CRISPR edits) using subsampled contact maps (details in Methods). (**D**) Results of CRISPR/Cas9-based deletion of an LTR41 element at chr8:70.3–71.8 Mb in GM187278 cells. Mega-contact maps generated in Hi-C^2^ experiments for the (top) WT locus and (bottom) KO (ΔLTR41) locus. (**E**) Virtual 4C plot displaying total percent interactions emanating from an anchor on a 5kb-window containing the left anchor CTCF of the conserved loop, and (**F**) the LTR41 element.

Our second candidate was a species-specific LTR41-derived CTCF site (“c” in Fig 3D, E) that replaced an ancestral CTCF site derived from a much older TE (“d” in Fig 3D, E) of the MER82 subfamily that is conserved in humans and mouse. The ancestral MER82-derived CTCF site was “decommissioned” as the LTR41 insertion (after the primate-rodent split) provided a negative orientation CTCF motif upstream of the MER82 element. Based on the loop extrusion model, the LTR41-derived CTCF motif would be encountered before the MER82-derived CTCF site and hence the ancestral site is mostly decommissioned in present-day human genome as evidenced by the drastically reduced CTCF binding (Supplementary Figure 4B). In the WT contact map, we observed a bright focal enrichment corresponding to CTCF binding sites a-c suggesting a looping interaction. In contrast, there was little focal enrichment corresponding to a-d (Fig 3D, top row). Additionally, in the WT Virtual 4C track anchored on “a”, we observed a clear peak corresponding to LTR41 (“c”) suggesting an a-c loop (Fig 3E). Upon deletion of LTR41, the conserved loop’s anchor is offset to the MER82-derived CTCF site (“d”) downstream of the LTR41 as evidenced by the shift in the focal enrichment in the KO contact map (Fig 3D, bottom row) and an increase in the KO Virtual 4C peak corresponding to the MER82-derived CTCF site (i.e., a-d loop) (Fig 3E, Supplementary Figure 7). Upon anchoring the Virtual 4C on a 5-kb window containing LTR41 (c), we observed a peak loss at “a” corresponding to the loss of the a-c loop in the KO, an interaction that existed in the WT cells (Fig 3F). With the ∼39kb shift of the anchor site, the half-megabase scale chromosomal structure around the anchor region remained largely preserved (Supplementary Figure 4C). Upon deletion of this TE candidate, the local sequence configuration probably resembled that of the pre TE-insertion, ancestral genome. This example therefore illustrates a potential path by which the local 3D genome evolved upon insertion of the LTR41 element as well as the plasticity TEs, like LTR41 and MER82 in this case, can encode in their host genomes by providing redundant CTCF binding sites.

These results support the hypothesis that TEs are able to contribute regulatory robustness and strengthen conserved regulatory architecture as redundant or “shadow” loop anchors. The mouse genome that underwent a lineage-specific expansion of SINE B2s (*22*), which carry a CTCF binding motif, is saturated with such events.

TEs are typically silenced by host repressive machineries including DNA and histone methylation (2*7-29*). However, a small fraction of TEs escape epigenetic silencing and provide functional regulatory elements for the host in a process termed exaptation (*30-33*). Since CTCF is a methylation sensitive chromatin factor and only binds to unmethylated DNA (*34,35*), we examined the DNA methylation levels of loop anchor CTCF sites of orthologous loops (Supplementary Methods). We found that TE-derived CTCF sites were marked by reduced DNA methylation, similar to their non-TE derived genomic counterparts (Fig 4A). To understand the DNA methylation dynamics through evolution, we took advantage of the differential mutation rate of 5-methylcytosine (5mC) to Thymine (T) (*36*). Unmethylated cytosines (C) mutate to T at a lower rate than 5mC; thus, methylated DNA exhibits higher frequency of C to T mutations (*37*). We found that TEs involved in turnover events had a significantly lower frequency of methylation-associated C-to-T and G-to-A mutations compared to an identically sampled background of TEs not involved in looping (1000 simulations), but no difference in all other combined substitutions (summarized human results: Fig 4B; full human and mouse results: Supplementary Figure 8, 9, Table S3). These results suggest that TEs providing CTCF turnover were hypomethylated over evolutionary time to maintain their functional role, compared to other TE copies (Fig 4C).

**Figure 4:**
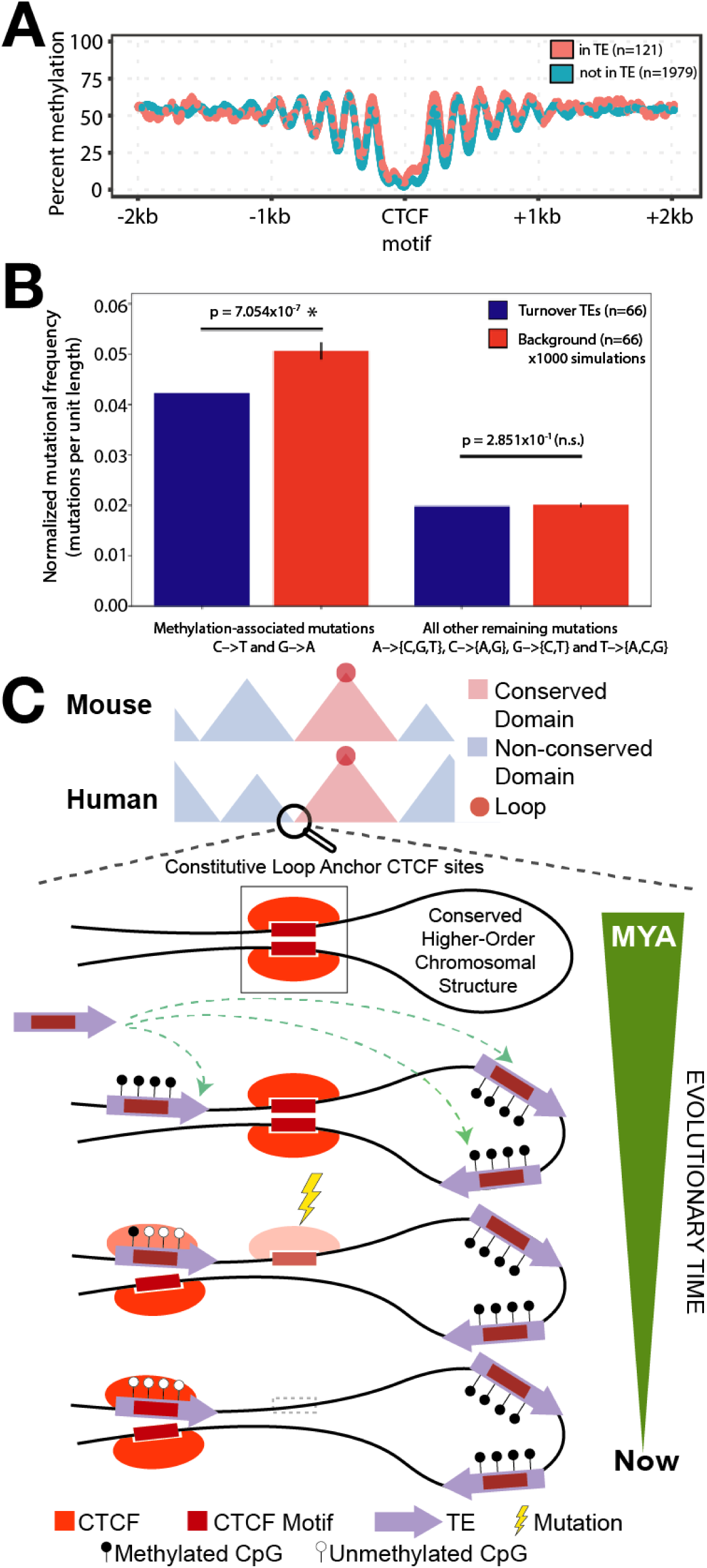
Turnover TEs are hypomethylated through evolutionary time. **(A)** Methylation signature ±2kb around CTCF sites that help maintain orthologous loops segmented by the origin of the anchor CTCF site (**B**) Methylation-associated and non-methylation mutational signature of individual TEs relative to its ancestral sequence in humans (mouse TE data available in Supplementary Figure 8). Alignments were performed using crossmatch (shown here) and Needle (details in Methods, results in Supplementary Figure 9). Error bars show one standard deviation of the means from 1000 simulations. (**C**) Schematic depicting the framework of TE-mediated CTCF binding site turnover that highlights the intimate reciprocity between the TE, genome and epigenome, to help maintain conserved 3D genome.

## DISCUSSION

TEs have substantially contributed to higher order chromatin structures by serving as chromatin loop anchors—a large fraction of which were found to be species-specific, confirming TEs’ role in genome innovation. Pioneering work in the last decade has extensively outlined this contribution of TEs in shaping gene regulatory networks by depositing TF binding sites in host genomes, leading to the origins of novel phenotypes like innate immunity and pregnancy in mammals. Herein, lies the catch: research to date showcases the role of TEs in bringing novelty and new regulatory functions to the host genome. Hence, TEs have long been considered a source of genetic innovation. However, by comparing topologies instead of raw DNA sequences in this study, for the first time, we have been able to reveal the role of TEs in 3D genome conservation. This seemingly counter-intuitive role of species-specific parasitic sequences in helping maintain ancestral genome architecture is fundamentally different from all current and previous work regarding TEs’ role in gene regulation. This role is mediated by a long-postulated, classic genetic phenomena of binding site turnover—for CTCF in this case. Redundant TE-derived CTCF sites in the vicinity of conserved chromatin anchor/ boundary can sometimes take over from the conserved anchor/boundary element, thus slightly shifting the anchor/boundary site while largely maintaining the 3D structure. Certain TE subfamilies like mouse SINE B2s contain pre-existing CTCF motifs within them, while others like mouse RLTR30 provide sequence fodder which upon a couple of specific point mutations can acquire CTCF binding and potentiate this binding site turnover.

In this study, 123 turnover events were observed in mouse on the basis of 3331 annotated loops (3.7%) whereas in humans 89 turnover events were observed out of 9448 loops (0.94%). This 4-fold higher rate of turnover events in mouse highlights differences in between species and the turnover phenomenon being investigated. The higher rate of loop anchor CTCF turnover in the mouse genome was amplified by the arrival of CTCF-motif containing B2 elements. The genome is replete with such events and we have for the first time functionally dissected and validated them in the context of 3D genome conservation, opening the doors up for such investigations in the field for enhancer or promoter turnover events.

The *fons et origo* of CTCF motifs in B2 SINEs has been extensively researched. B2 SINEs are derived from tRNA genes. Mouse tRNA genes have been shown to possess classical insulator activity and the potential to function as boundary elements (*38*). Moreover, CTCF-binding enrichment in B2 SINEs and repeat-driven dispersal of CTCF-binding has been shown to be a fundamental, ancient, and still highly active mechanism of genome evolution in mammalian lineages (*22*).

Similarly, the role CTCF motifs in viral genome regulation has been a topic of tremendous interest and investigation. In EBV, this control involves direct binding of CTCF across the viral genome and the formation of three-dimensional loops between virus promoters and enhancers (*39*). CTCF has been shown to be important in the regulation of gene expression of a number of human DNA viruses (*40*). CTCF also plays a critical role in epigenetic regulation of viral gene expression to establish and/or maintain a form of latent infection that can reactivate efficiently (*41*). Recent evidence has also shown that HTLV-1 inserts an ectopic CTCF binding site forming loops between the provirus and host genome, altering expression of proviral and host genes (*42*). CTCF has also been shown to promote HSV-1 lytic transcription by facilitating the elongation of RNA Pol II and preventing silenced chromatin on the viral genome (*43*). Moreover, one can speculate that having a CTCF motif can not only help in maintaining viral genome confirmation but can also help insulate the chromatin activity of the neighborhood wherein the virus inserts into the host genome. It may also increase the chances of long-range interactions taking place which can sometimes bring in other TFs and/or polymerase, leading to enhanced transcription at the site of viral integration.

Our in-depth analysis of 3D genome structures upon genetic manipulation of candidate TEs revealed principles of how 3D genome evolves. In one example, a human TE provided a conserved chromatin boundary and loop anchor, whereas the ancestral CTCF site had decayed. Upon deletion, the chromatin domains collapsed, and loops eliminated, underscoring the importance of the TE in maintaining the local 3D genome structure.

In another case where a human TE provided a similarly conserved boundary and loop anchor, the ancestral CTCF site was still recognizable but was decommissioned. Deletion of the TE resulted in reinstallation of the ancestral CTCF site to form a slightly shifted boundary and loop anchor, and the local chromatin domains were largely preserved. In this second case that we validated, we undid the events that took place during the course of (tens of millions of years) evolution by removing a young TE (LTR41) and having the ancestral “decommissioned” TE (MER82) re-uptake its function, thereby “reversing” the path of evolution in a dish (in days). Thus, experimentally demonstrating the evolutionary impact of a TE-derived CTCF site. Moreover, the concept of such shadow loop anchors residing in TEs that can be activated upon escape from epigenetic silencing is extremely crucial to take into account for studies pertaining to diseases of the epigenome like certain cancers, their treatment and therapy. This study also underscores the redundancy that exists in the genome when it comes to CTCF binding sites and can potentially explain why we may not always see a change in 3D genome structure upon deleting CTCF binding sites.

It is important to remember that the contribution outlined in this manuscript are underestimates as we have yet to (i) uniquely identify all loop anchor CTCF sites (especially in highly repetitive regions), (ii) annotate all repetitive elements, especially ancient TEs that have diverged far from their identity (*18*), and (iii) identify other architectural proteins and expand this framework beyond just CTCF-derived loop anchors.

While most studies highlight TEs’ role in innovating new functions by providing novel regulatory elements such as enhancers and promoters, we implicate the role of TEs in functional conservation inviting us to reexamine this unconventional role—perhaps many novel regulatory elements derived from TEs are not creating new functions, but rather providing redundant genetic material thus contributing to the robustness of gene regulatory networks. These findings will undoubtedly stimulate investigations to explore the multitude modes of regulatory evolution mediated by TEs. Indeed, recent evidence has linked the transcriptional activation of retrotransposons to restructuring of genome architecture during human cardiomyocyte development (*44*).

A major caveat of the analysis presented in this study is that the *in situ* Hi-C maps (re-analyzed in this study) of the 9 cell lines were sequenced to varying depths, and thus differ in their resolution and “completeness” of loop annotations. Hence, due to this limitation of publicly available high-resolution HiC data, our findings likely represent a lower bound of TE’s involvement in shaping both the conserved and species-specific 3D genome. These analyses need to be revisited as and when higher-resolution datasets are available.

Lastly, our study opens the doors for population-scale genetic variation studies that identify polymorphic TE insertions to be reconciled with population-scale 3D genome and regulatory variation. These future explorations will present yet another vignette of transposable elements and their very many roles in accelerating adaptive evolution.

## CONCLUSIONS

Taken together, our findings reveal a formerly uncharacterized role that TEs have played in the evolution of higher-order chromosomal structures in mammals. TEs have contributed a substantial number of loop anchors in mouse and human 3D genomes, a fraction of which were co-opted to help maintain conserved higher-order chromosomal structures. TE transposition provides redundant CTCF motifs and a novel method for CTCF binding site turnover to maintain regulatory conservation (defined here as the preservation of long-range chromosomal interactions, loop and boundary formation), by compensating for the loss of local primary sequence—local sequence that would have otherwise allowed the assessment of purifying selection. Deletion of these TEs in human cell lines eliminated the chromatin loops that they anchor and resulted in collapse of conserved chromatin structure, as expected by our hypothesis. More strikingly, we demonstrate that in another case the loop anchor shifted to an alternative TE-derived CTCF site nearby, resulting in largely unchanged chromatin structure, underscoring the dynamic nature and robustness of the 3D genome upon TE infiltration. These TEs that maintain conserved chromatin loops via turnover are hypomethylated through deep time, an observation that highlights the intimate interplay between genome, epigenome, and 3D genome in evolution. This research provides a foundation to study the impact of TEs and expand our understanding of chromosomal folding—its emergence, maintenance and transformation—in the context of evolving genomes. Ultimately, our study reveals how selfish genetic elements, regardless of their origins, can be repurposed to provide redundant TF motifs, maintain latent genome sanctity and regulatory fidelity by conserving 3D structure.

## METHODS

### Dataset GEO accession numbers

The genomic data analyzed in this study were obtained from publicly available datasets. HiC datasets were obtained from GSE63525 (mouse: CH12; humans: GM12878, HeLa, HMEC, IMR90, K562, NHEK). GM12878 ChIA-PET dataset was obtained from GSE72816. GM12878 CTCF ChIP-seq datasets were obtained from ENCODE (ENCSR000AKB and ENCSR000DZN). CH12 CTCF ChIP-seq datasets were obtained from Mouse ENCODE (ENCSR000ERM and ENCSR000DIU). WGBS methylation dataset for GM12878 was also obtained from ENCODE, GEO: GSE86765 (ENCSR890UQO). Mouse ESC and NSC HiC data was obtained from PMID: 30414923.

### Loop anchor CTCF–RE intersection

We generated a list of unique anchor CTCF sites using the HiCCUPS output^1^ for various mentioned cell lines. We then overlapped loop anchor CTCF motifs identified using HiCCUPS with *RepeatMasker* (RMSK v4.0.7, for hg19 and mm9) and required at least 10bp of the core CTCF motif to intersect with a repetitive element (RE) to call it a RE-derived loop anchor CTCF site. Further, only loops with (i) at least one known RE-derived anchor CTCF site, or (ii) two non-RE derived anchor CTCF sites were taken into consideration for analysis of RE-derived loop counts, because we can definitively say whether the loops and their loop anchor CTCF sites were derived from REs or not. Loops with both unidentified loop anchor CTCF sites, or one unidentified and one non-RE derived anchor CTCF site were not considered as there is the possibility of having at least one of the other anchor CTCF sites derived from a RE. We followed the same methodology when considering ChIA-PET loops.

### TE class and family distribution

We ran RepeatMasker v4.0.7 with the -s slow search parameter on the hg19 and mm9 genomes to obtain a comprehensive list of REs in the genome and their corresponding subfamily, family and class annotations. We used RE counts (generated as previously outlined) to characterize their distribution to loop anchor CTCF sites. For characterizing RE-derived CTCF binding peaks, we repurposed a previously used strategy (*10*). Briefly, we required that the centers of the MACS-called peaks of ENCODE-generated CTCF ChIP datasets overlapped with RE fragments. We used the length distribution of various RE family and classes in the entire genome as the background distribution.

### Loop orthology check

We used liftOver (*19*) to convert CH12 loop annotations from mm9 mouse genome coordinates to hg19 human genome coordinates. We used various sequence match rates (minMatch = 0.05, 0.1…, 1) to convert CH12 mouse peaks from mm9 genome coordinates to hg19 genome coordinates. To optimize for the minMatch parameter, we generated ten shuffled (randomized) peak annotations by using bedtools shuffle –chrom command to permute their location on the chromosome of origin. minMatch parameter of 0.1 was chosen for liftOver analyses henceforth, as it resulted in the greatest number of features being lifted over (on average) and lower coefficient of variation across the 10 simulated sets. We lifted over 3245 out of 3331 mouse peaks from mm9 to hg19, using the minMatch 0.1, to facilitate cross-species peak annotations comparison. To call a mouse feature conserved in humans, we required that the loop anchor pairs individually lie within a min(half of loop length, vicinity threshold) window of an existing loop anchor pair. The vicinity threshold was put in place to account for cross-species liftOver errors and facilitate comparison of higher-order chromosomal features that vary from 120Kb to 125Mb in length (in mouse). We tested multiple vicinity thresholds ranging from 500bp to 100Mb and identified false discovery rates using simulated sets of mouse features and comparing them to the orthology observed between the real CH12 (mouse) and GM12878 (human) features. We decided to use 50kb as the vicinity threshold as it corresponded to a false discovery less than 0.1. We found that 1688 CH12 mouse peaks overlapped at least one corresponding peak in GM12878 human lymphoblastoid cells. We performed a similar analysis to compare ‘muranized’ human features (liftOver from GM12878) to actual mouse features (CH12). We found that 1900 GM12878 human peaks overlapped at least one corresponding peak in CH12 mouse lymphoblastoid cells. We then filtered for features that displayed reciprocal matches (reciprocal best hits) in the two comparisons (mouse-to-human and human-to-mouse) as stated above. Finally, we curated the list by considering genic, epigenomic and transcriptomic synteny to pick exactly one orthologous human loop to a corresponding mouse loop, to enlist 1596 high-confidence orthologous peak calls (Table S1.1). A brief flowchart of the pipeline is shown below:

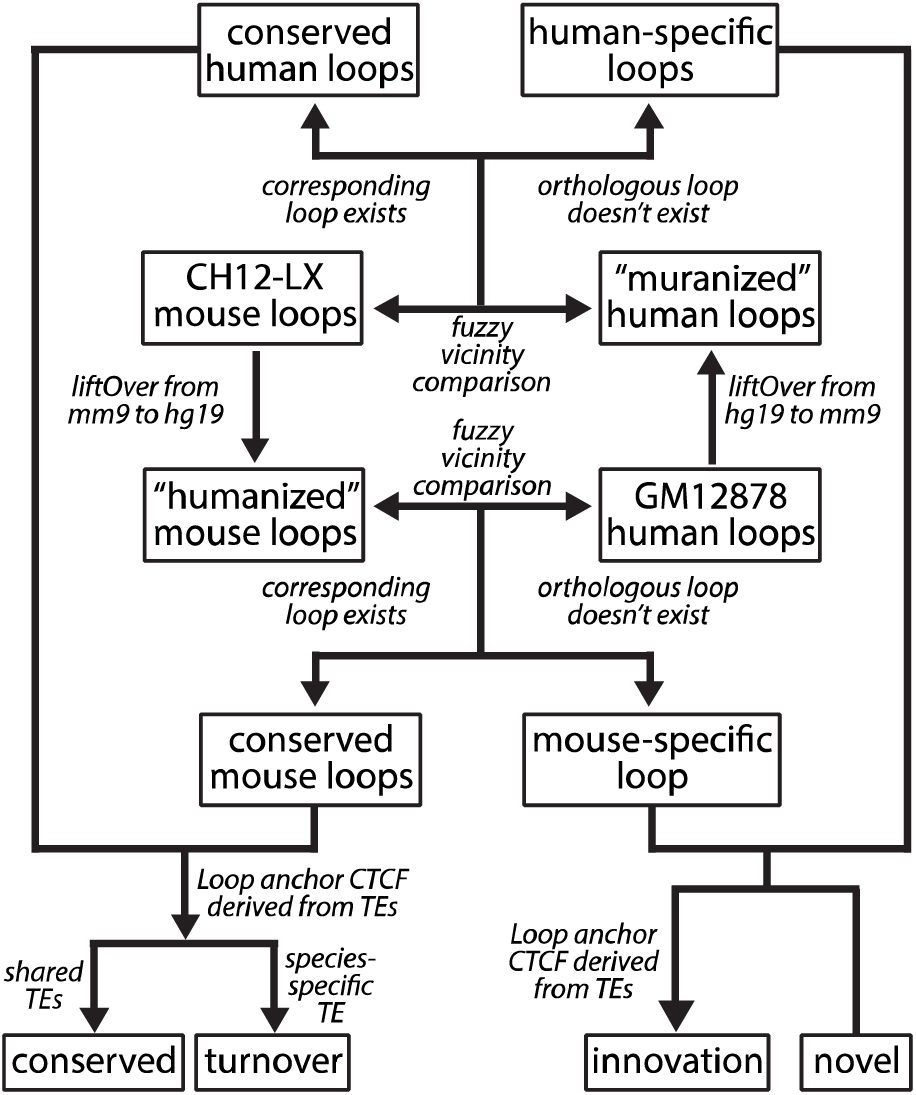

### TE age estimation

Species divergence times were based on (*45*). Repeat ages were estimated by dividing the percent divergence of extant copies from the consensus sequence by the species neutral substitution rate. Substitution rates (mutations/yr) used were as follows: humans: 2.2×10^−9^; mouse: 4.5×10^−9^, from (*46*). Jukes-Cantor and Kimura distances were calculated by aligning each TE to its consensus sequence and counting all possible mutations (see below). Single nucleotide substitution counts were normalized by the length of the genomic TE minus the number of insertions (gaps in the consensus). These mutation rates were then used to calculate the Jukes-Cantor and Kimura distances for each genomic TE.

### Candidate selection and filtering

After manually curating the list of conserved loops, we looked for TE-derived orthologous loops in humans that were discordant for TEs in mouse. After identifying the list of TE-derived CTCF turnover events in humans, we comprehensively surveyed the local CTCF binding landscape (CTCF ChIP-seq peaks) to ensure (i) there weren’t other CTCF binding sites in the vicinity that could function as loop anchors in humans (in the first case); and (ii) there was only one other unique CTCF binding site, i.e. the ancestral CTCF motif (in the second case). We also ensured that the TE insertion from which the loop anchor CTCF site was derived was human-specific and not present in mouse (Table S1.2). We repeated this analysis to identify TE-mediated turnover in mouse as well (Table S1.3). We also identified events wherein TEs mediated turnover events both in mouse as well as human (Table S1.4). One possible explanation for this observation is that similar selective pressures (i.e. the need to maintain higher-order chromosomal structure) led to the convergent co-option of species-specific TEs at syntenic locus, independently in both the genomes.

### Cell culture methods

GM12878 cell lines were grown between 200K-800K cells/ml in 10ml cultures in T-25 flasks, in a humidified incubator with 95% CO_2_ at 37°C in RPMI1640 media (Gibco, 1187-085) supplemented with 15% fetal bovine serum (Corning, 35-011-CV) and 100U/ml penicillin-streptomycin (Gibco, 15140-122) as per the ENCODE standards.

### CRISPR-Cas9 mediated genome engineering

Our CRISPR workflow consisted of the following steps: We identified turned over chromatin loops that are maintained by TEs, with unique, correctly oriented TE-derived CTCF motifs within loop anchors (*1*). We used two independent CRISPR sgRNA design engines CRISPOR (*47*) and CRISPRScan (*48*) to rationally design multiple pairs of sgRNAs that have high cutting efficiency and minimizing off-target effects. We used pU6-(BbsI)_CBh-Cas9-T2A-BFP plasmid (Addgene, 64323) and pU6-(BbsI)_CBh-Cas9-T2A-mCherry plasmid (Addgene, 64324) as the CRISPR delivery vectors. For each sgRNA, we designed and annealed two single-stranded oligos with compatible overhangs that can be cloned into BbsI-digested BFP and mCherry CRISPR vectors through standard ligation techniques. For every pair of sgRNAs, we constructed BFP-CRISPR vectors and mCherry-CRISPR vectors that express sgRNAs targeting upstream and downstream of the candidate TEs, respectively. BFP-CRISPR vectors and mCherry-CRISPR vectors each were co-transfected into GM12878 cells in antibiotic-free media using the Neon transfection system. After 24 hours of incubation, the transfected cells were analyzed by flow-cytometry (Beckman Coulter MoFlo) for BFP-positive and mCherry-positive subpopulations. Transfection efficiencies were usually between 3-5%. We single-cell sorted these double-positive fluorescent cells into 96-well plates for clone expansion and allowed to grow for 21-28 days. After that, 20-48 clones were screened per transfection. Genomic DNA from CRISPR clones was extracted using *Quick-DNA* Miniprep kit for genotyping and validated with Sanger sequencing. Details of sequences used to generate clones used in this study are listed in Table S2.4. We then performed *in situ* Hi-C on the selected mutated cell lines and performed hybrid selection on the in situ Hi-C libraries for a region around the targeted TE to generate Hi-C^2^ libraries that can easily and cheaply be sequenced to read off the effects of our TE deletions on local genome folding.

### Hi-C^2^ probe design

To design probes targeting the two regions for HYbrid Capture Hi-C (Hi-C^2^), we followed a similar approach as (*8*). In short, we (i) identified all MboI restriction sites within the target region, (ii) we designed our bait probe sequences to target sequences within a certain distance of the MboI restriction sites as Hi-C ligation junctions occur between them, (iii) we followed a similar three-pass probe design strategy sequentially increasing various parameters like the distance of the probe from the MboI restriction site, the number of repetitive bases, the GC content, probe density in gaps with relaxed probe design quality filters. We then removed overlapping probes or probes with identical sequences. After all three passes, we identified 2741 unique probes covering region 1 (chr10:26-28Mb; 1.37 probes/kb) and 1856 probes covering region 2 (chr8:70.3-71.8Mb; 1.24 probes/kb). 15bp primer sequences (unique for each region, details in Table S2.3) were then appended to both ends of the 120bp probe sequence to facilitate single oligo pool synthesis and subsequent amplification of region-specific sub-pools. Probe construction and hybrid selection was then followed with sequences specific to this study using the same strategy detailed in (*8*).

### Hi-C experiments

The Hi-C datasets used in our analyses were generated using the *in situ* Hi-C protocol standardized by the 4DN consortia. In brief, the in situ Hi-C protocol involves crosslinking cells with 1% formaldehyde for 10 minutes, permeabilizing them with nuclei intact, digesting the DNA with MboI (4-cutter restriction enzyme), filling the 5’-overhangs while incorporating biotin-14-dATP (a biotinylated nucleotide), followed by ligating the resulting blunt-end fragments, shearing the DNA to a 400-700bp fragment size, capturing the biotinylated ligation junctions with streptavidin beads, building an Illumina library with 10-12 rounds of PCR amplification, and finally analyzing the resulting fragments with paired-end sequencing. The resulting library was always shallow sequenced to 500K-4M reads to check for library build quality looking at key statistics such as complexity, number of Hi-C contacts, inter vs. intrachromosomal interactions, and long-range vs/ short-range intrachromosomal interactions. Libraries that passed the quality check were either sequenced deeper and/or used as pools for subsequent Hi-C^2^ experiments.

For our genome engineering experiments, we generated 14 in situ Hi-C libraries (Table S2.1) from GM12878 cells. We also generated 16 *in situ* Hi-C^2^ libraries from various genome-engineered GM12878 cell lines on which we performed hybrid selection. All *in situ* Hi-C libraries generated as part of this study are detailed in Table S2.2. All the Hi-C data was processed using the computational pipeline described in full detail in (*1*). Hi-C libraries were sequenced to a depth of between 624K-333M reads (on average, 63.8M reads). Hi-C^2^ libraries were sequenced to a depth of between 6.7M-168M reads (on average, 35.8M reads). All data was initially processed using the pipeline published in (*1*) and visualized on the desktop and web version of Juicebox. We combined Hi-C and Hi-C^2^ contact maps corresponding to the same genotype and the same locus using the Juicer’s mega.sh script as these are in essence “biological” replicates, to generate higher resolution megamaps.

### Analysis of cross-domain interactions

We subsampled the Hi-C^2^ corresponding to the R1-WT megamap (containing 46M reads) and R1-KO (containing 56M reads) for 5M reads, 10 times to create 10 independent R1-WT and R1-KO mini-maps. For each of these HiC maps, we used the Juicer Tools dump command to extract the raw contact matrix. Intradomain interactions were defined as interaction that (i) originate and terminate in domain 1, or (ii) originate and terminate in domain 2. Interdomain interactions were defined as interactions that originate in domain 1 and terminate in domain 2. We then calculate percentage of cross-domain interactions for each of the mini-maps using the formula: (number of intradomain-interactions)*100 / ((number of intradomain-interactions) + (number of interdomain-interactions)). The percentage of cross-domain interactions were calculated for the target domain as well as a control domain. The distribution of cross-domain interactions across the targeted domain was found to be significantly different in the KO vs. the WT (t-Test: Two-Sample Assuming Unequal Variances, p-value = 1.40668×10^−16^). The distribution of cross-domain interactions across a nearby control domain however was not found to be significantly different in the KO vs. the WT (t-Test: Two-Sample Assuming Unequal Variances, p-value = 0.013254165). Raw simulation data and statistics are provided in Table S2.5.

For Figure S3C, we used the Hi-C megamap corresponding to R2-WT and R2-KO to retrieve raw interaction counts at a 100kb resolution. Percent cross-domain interactions was calculated using the formula stated above. We calculated the enrichment of cross-domain interactions in the LTR41-DKO w.r.t. the WT across the targeted domain as well as a nearby control domain.

### DNA Methylation analysis

We generated a methylation metaplot representing the mean CpG methylation value from WGBS data (ENCODE dataset: ENCFF835NTC) of 20bp sliding windows, centered on CTCF motifs (and ±2kb around it) segmented by their origin/TE-derivation status.

### Analysis of TE Mutational Profile

#### 1. TE consensus construction

For most of the TE subfamilies, we retrieved the consensus sequences from the RepBase library (RepBase 22.02, RepeatMaskerEdition20170127) (*49*). However, LINE elements are fragmented to 5’ end, ORF2 and 3’ end regions in RepBase library. To reconstruct full-length LINE consensus, we identified TE fragments in human and mouse genome using RepeatMasker and compared the standard output (.out file) with the alignment output (.align file) from the same RepeatMasker run (*50*). For each LINE element in the standard output, we summarized which 5’ end, ORF2, and 3’ end fragments have been used most to construct the full-length element. Then we use EMBOSS Water local alignment algorithm to align the three pieces together and generated the full-length LINE consensus sequences (*51*).

#### 2. Crossmatch alignments

We ran RepeatMasker 4.0.7 on the mm9 and hg19 genomes using crossmatch as the search engine. We then parsed the alignment file to determine the substitution rates between the ancestral sequence and the genomic element. For each genomic element, we counted the number of A-to-C, A-to-G, A-to-T, C-to-A, C-to-G, C-to-T, G-to-A, G-to-C, G-to-T, T-to-A, T-to-C, and T-to-G substitutions (single nucleotide substitutions), where the first nucleotide indicates the ancestral sequence and the second nucleotide indicates the genomic sequence. We ignored any substitutions that involved ambiguous nucleotides. We also counted the number of insertions and deletions. All substitution frequencies were normalized by the length of the genomic sequence to estimate the substitution rates in each TE. Any genomic TE with a length less than 20% of the ancestral sequence was filtered out. For each single nucleotide substitution, we calculated the average substitution rate in two subsets of TEs (details below). We also calculated the combined C-to-T and G-to-A substitution rate (methylation-associated substitutions) and the combined rate of all other substitutions (non-methylation-associated substitutions) to compare the rate of DNA methylation-induced mutations to other mutations. The methylation substitution rate was computed by taking the average of the C-to-T and G-to-A rates for each TE and then averaging over turnover events. The non-methylation substitution rate was computed by taking the average of all other (ten) single nucleotide substitutions for each TE and then averaging over turnover events.

We generated a background distribution by repeating this analysis on 1000 permutations of all genomic TEs. We first calculated the frequency of each TE subfamily in the set of turnover events. For each permutation, we randomly selected genomic TEs (not involved in anchoring loops) from each subfamily to reflect their frequency in turnover events. The single nucleotide substitution rate, methylation-associated substitution rate, and non-methylation-associated substitution rate were calculated as described above. The distribution of all substitution rates from the permutations follow a normal distribution (KS test, P > 0.0036, Bonferroni correction alpha = 0.05 for N = 14 hypotheses, Supplemental Table S3.1). The background distribution was then used to perform a left-tailed z-test. We did not compute a two-tailed p-value because our null hypothesis is that the observed mutation rates are greater than or equal to the background distribution mean. For the 12 single nucleotide substitutions, we used Bonferroni correction to account for multiple hypotheses.

#### 3. Needle realignments

RepeatMasker performs post-processing after running crossmatch, so coordinates and TE subfamily assignments in the .out file do not always reflect the contents of the .align file. To improve our estimates of mutation rates, we realigned each TE to its matched consensus sequences. We extracted the genomic and subfamily consensus sequence using the coordinates reported in the .out file. We then performed a global alignment using EMBOSS Needle v6.6.0.0 using a gap open penalty of 10, a gap extension penalty of 0.5, and the EDNAFULL scoring matrix. We used the alignment to recompute single nucleotide substitutions for each TE and then repeated the same analysis we used for crossmatch alignments. We did not filter out TEs with a length less than 20% of the ancestral sequence because this filter was originally put in place to account for discrepancies between the .align and .out files. As before, the distribution of all substitution rates from the permutations follow a normal distribution (KS test, P > 0.0036, Bonferroni correction alpha = 0.05 for N = 14 hypotheses, Supplemental Table S3.2).

## DECLARATIONS

### Ethics approval and consent to participate

Not applicable

### Consent for publication

Not applicable.

### Availability of data and material

The data sets generated and analyzed in this current study will be uploaded to GEO upon acceptance

### Authors’ Contributions

M.N.K.C. and T.W. conceived and designed this study; M.N.K.C. analyzed the data, performed experiments, generated sequencing libraries, and wrote the manuscript with inputs from T.W.; R.Z.F., J.T.W., and X.Z. contributed text and revised the manuscript; H.S.J. contributed reagents and resources; R.Z.F performed mutation frequency simulations along with M.N.K.C.; X.Z. generated TE ancestral sequences and TE alignments; T.W. supervised the project. All authors subsequently edited and approved the final manuscript.

### Supplementary Materials

Table S1 – S3

Supplementary Figure 1 – 9

## Acknowledgements

We thank members of the Wang Lab for helpful discussions related to the project; Jessica Hoisington-López and Maria Lynn Jaeger from The Edison Family Center for Genome Sciences & Systems Biology for assistance with sequencing; Matthew Patana & Daniel Schweppe from the Siteman Flow Cytometry core for FACS expertise.

## REFERENCES

1. SSP Rao et al., Cell 159, 1665–1680 (2014).

2. Z Tang et al., Cell 163, 1611–1627 (2015).

3. T Sexton et al., Cell 148, 458–472 (2012).

4. JR Dixon et al., Nature. 485, 376–380 (2012).

5. DU Gorkin et al., Cell Stem Cell 14, 771–775 (2014).

6. W Schwarzer et al., Nature 551, 51–56 (2017).

7. AR Strom et al., Nature 547, 241–245 (2017).

8. AL Sanborn et al., Proc Natl Acad Sci. 112, E6456–E6465. (2015).

9. G Fudenberg et al., Cell Rep. 15, 2038–2049 (2016).

10. V Sundaram et al., Genome Res. 24, 1963–1976 (2014).

11. G Bourque et al., Genome Res. 18, 1752–1762 (2008).

12. G Kunarso et al. Nat Genet. 42, 631–634 (2010).

13. PÉ Jacques et al., PLoS Genet. 9(5) (2013).

14. T Wang et al., Proc Natl Acad Sci. 104, 18613–18618 (2007).

15. EB Chuong et al., Science 351, 1083–1087 (2016).

16. VJ Lynch et al., Nat Genet 43, 1154–1159 (2011).

17. BJ Britten, EH Davidson. Q Rev Biol. 46, 111–138 (1971).

18. APJ de Koning et al., PLoS Genet. 7 (2011).

19. AS Hinrichs, Nucleic Acids Res. 34, 590–598 (2006).

20. C Feschotte, EJ Pritham. Annu Rev Genet. 41, 331–368 (2007).

21. V Sundaram, T Wang. BioEssays. 40, 1700155 (2018).

22. D Schmidt et al., Cell 148, 335–348 (2012).

23. MZ Ludwig et al., Development 12, 3325–3330 (1998).

24. AM Moses et al., PLoS Comput Biol. 2, 130 (2006).

25. S Venkataram, JC Fay. Genome Biol Evol. 2, 851–858 (2010).

26. D Villar et al., Nat Rev Genet. 15, 221–233 (2014).

27. MA Matzke et al., Plant Mol Biol. 43, 401–415 (2000).

28. JA Yoder et al., Trends Genet. 13, 335–340 (1997).

29. RK Slotkin et al., Nat Rev Genet. 8, 272–285 (2007).

30. A Huda et al., Mob DNA 1, 2–12 (2010).

31. CB Lowe, D Haussler. PLoS One 7, 43128 (2012).

32. G Bejerano et al., Nature 441, 87–90 (2006).

33. MG Kidwell, DR Lisch. Trends Ecol Evol. 15, 95–99 (2000).

34. C Kanduri et al., Curr Biol. 10, 853–856 (2000).

35. S Kurukuti et al., Proc Natl Acad Sci. 103, 10684–10689 (2006).

36. JC Shen et al., Nucleic Acids Res. 22, 972–976 (1994).

37. AP Bird. Nucleic Acids Res. 8, 1499–1504 (1980).

38. T Ebersole et al. Cell Cycle. 10, 2779–2791 (2011).

39. I Tempera et al. PLoS Pathog 7:e1002180 (2011).

40. I Pentland et al. Viruses 7:3574–85 (2015).

41. JS Lee et al. mBio 9:e02372–17 (2018).

42. Y Satou et al. Proc Natl Acad Sci. 113(11):3054–3059 (2016).

43. F Lang et al. Sci. Rep. 7, 39861 (2017).

44. Y Zhang, T Li, S Preissl et al., biorXiv. (2018).

45. RW Meredith et al., Science 334, 521–524 (2011).

46. RH Waterston et al., Nature 420, 520–562 (2002).

47. M Haeussler et al., Genome Biol. 17, 148 (2016).

48. MA Moreno-Mateos et al., Nature Methods 12, 982–988 (2015).

49. W Bao et al., Mob DNA 6, 11 (2015).

50. A Smit et al., RepeatMasker Open-4.0.6 2013-2015. (2017).

51. P Rice et al., Trends Genet. 16, 276–277 (2000).

