## Supplementary Figures for "Co-opted transposons help perpetuate conserved higher-order chromosomal structures"

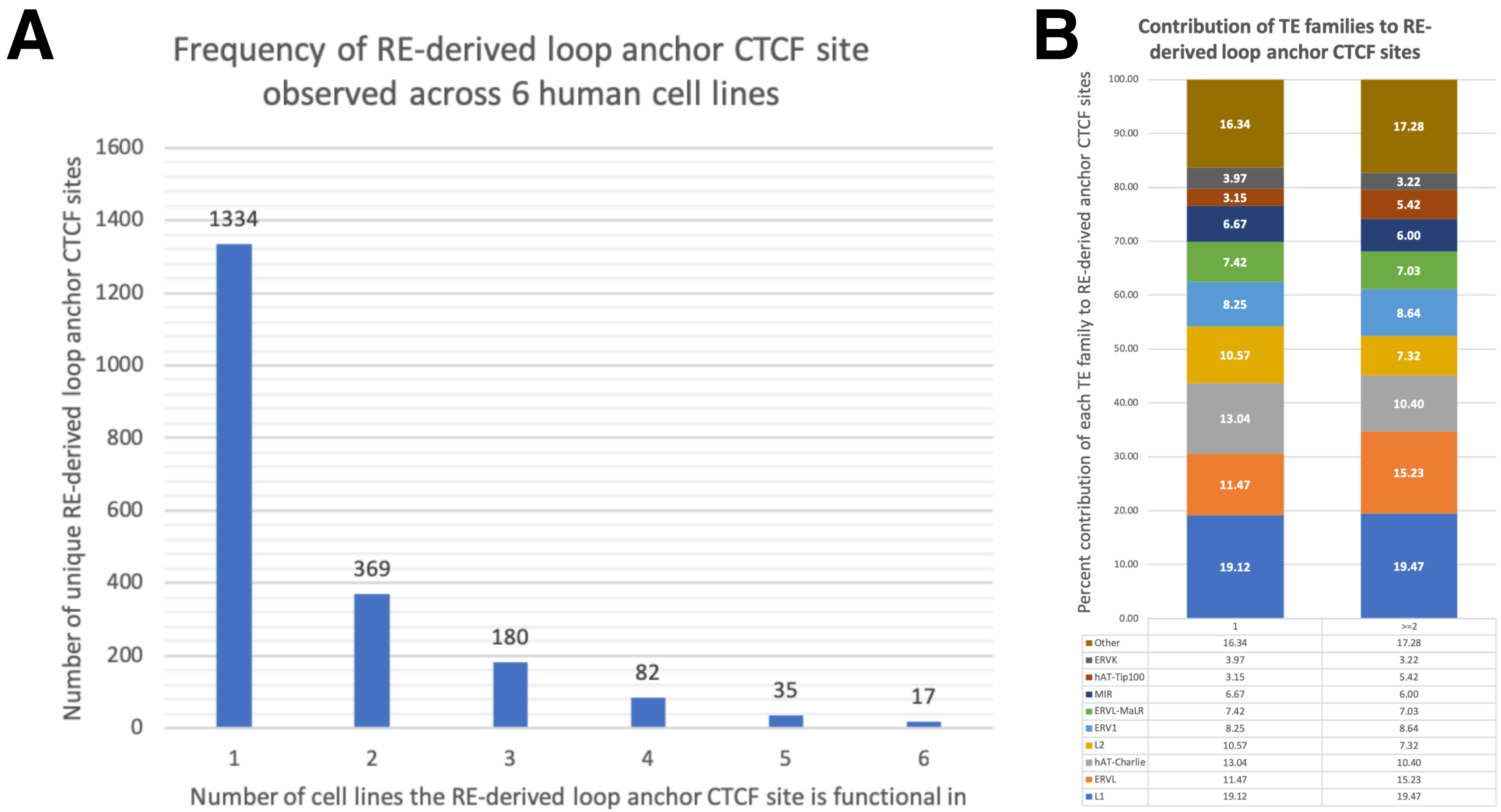


**Supplementary Figure 1 | Contribution of repetitive element (RE) families to cell-type specific loop anchor CTCF sites.** (**A**) Bar plot displaying the number of human cell lines in which a certain RE-derived CTCF site is a loop anchor. (**B**) Contribution of TE families to RE-derived loop anchor CTCF sites segmented by cell-type specific (found in exactly one sample) loop anchors (n=1334) and non-cell type specific (found in >=2 samples) loop anchors (n=683). (Note only top 9 TE families are displayed)

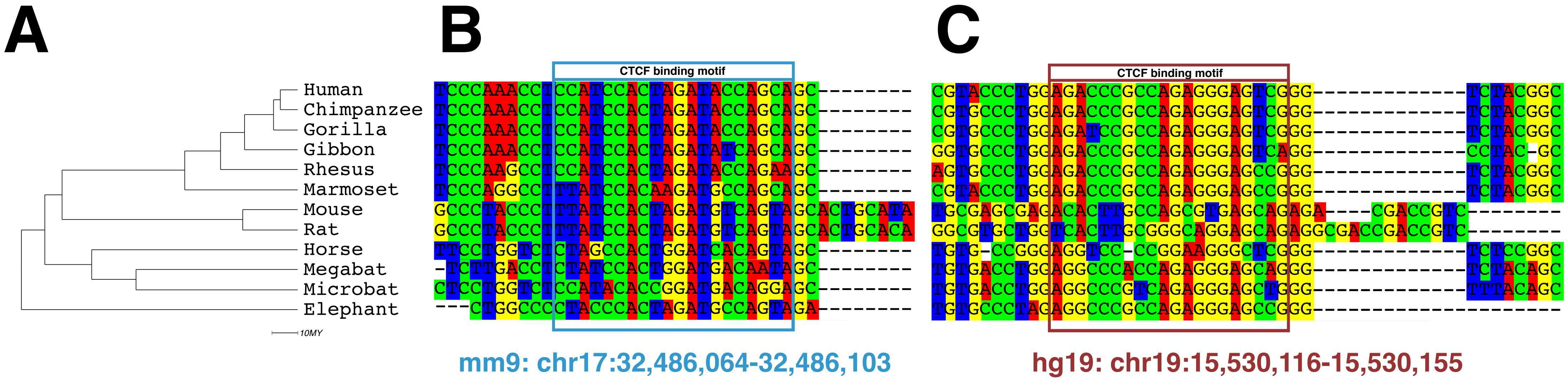


**Supplementary Figure 2 | Phylogenetic reconstruction of CTCF binding motifs that anchor the conserved chromatin loop near AKAP8L gene.** (**A**) Phylogenetic tree of various mammals from human to elephant. MY, million years ago. Representative species for each lineage is: *Homo sapiens* (human), *Pan troglodytes* (chimpanzee), *Gorilla gorilla gorilla* (gorilla), *Nomascus leucogenys* (Gibbon), *Macaca mulatta* (Rhesus), *Callithrix jacchus* (marmoset), *Mus musculus* (mouse), *Rattus norvegicus* (rat), *Equus caballus* (horse), *Pteropus vampires* (megabat), *Myotis davidii* (microbat), *Luxodonta africana* (elephant). (**B**) Syntenic sequences from various species corresponding to a 40bp region, that contains the turned-over CTCF binding motif (highlighted in blue box) derived from a MER20B element. (**C**) Syntenic sequences from various species corresponding to a 40bp region, that contains the ancestral CTCF binding motif (highlighted in red box) preserved in most non-rodent mammals.

*1) Xie X, Mikkelsen TS, Gnirke A, Lindblad-Toh K, Kellis M, Lander ES. Systematic discovery of regulatory motifs in conserved regions of the human genome, including thousands of CTCF insulator sites. Proc Natl Acad Sci USA. 2007;104:7145–50.*
*2) Ziebarth JD, Bhattacharya A, Cui Y (2013) CTCFBSDB 2.0: a database for CTCF-binding sites and genome organization. Nucleic Acids Research. 41(D1):D188-D194.*

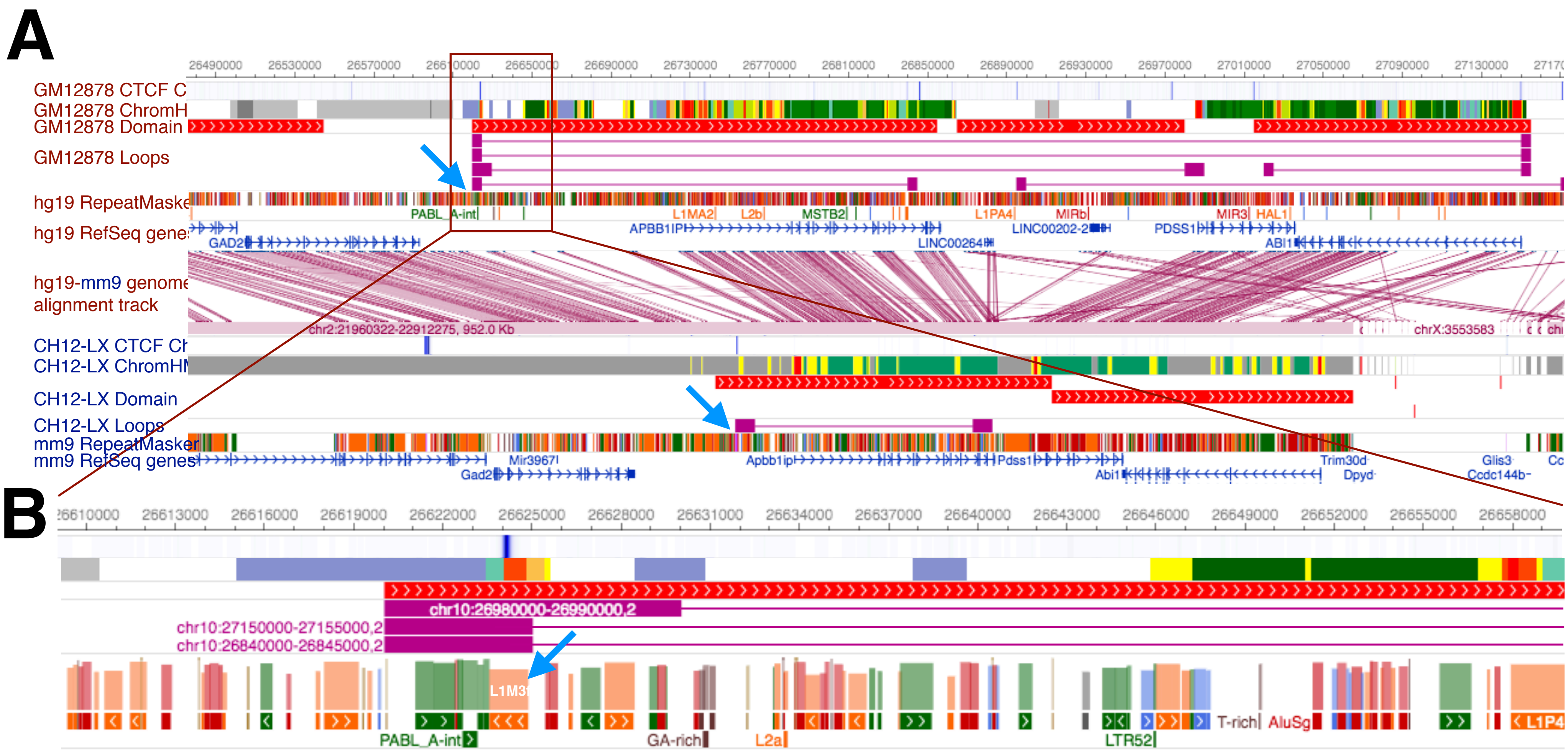


**Supplementary Figure 3 | Landscape of the higher-order chromosomal structures conserved in human and mouse maintained by an L1M3f-derived CTCF binding site turnover event in humans.** (**A**) Epigenome browser screenshot displaying the conserved genomic and epigenomic landscape. The conserved domain border and loop anchor in human and mouse is marked by blue arrows. Exactly one CTCF ChIP peak is observed in the vicinity of the domain border and loop anchor. (**B**) Zoomed in view of the conserved domain and loop anchor in humans shows that CTCF is bound to an L1M3f element (marked by a blue arrow).

*Visualization from WashU Epigenome Browser: http://epigenomegateway.wustl.edu*

**A**

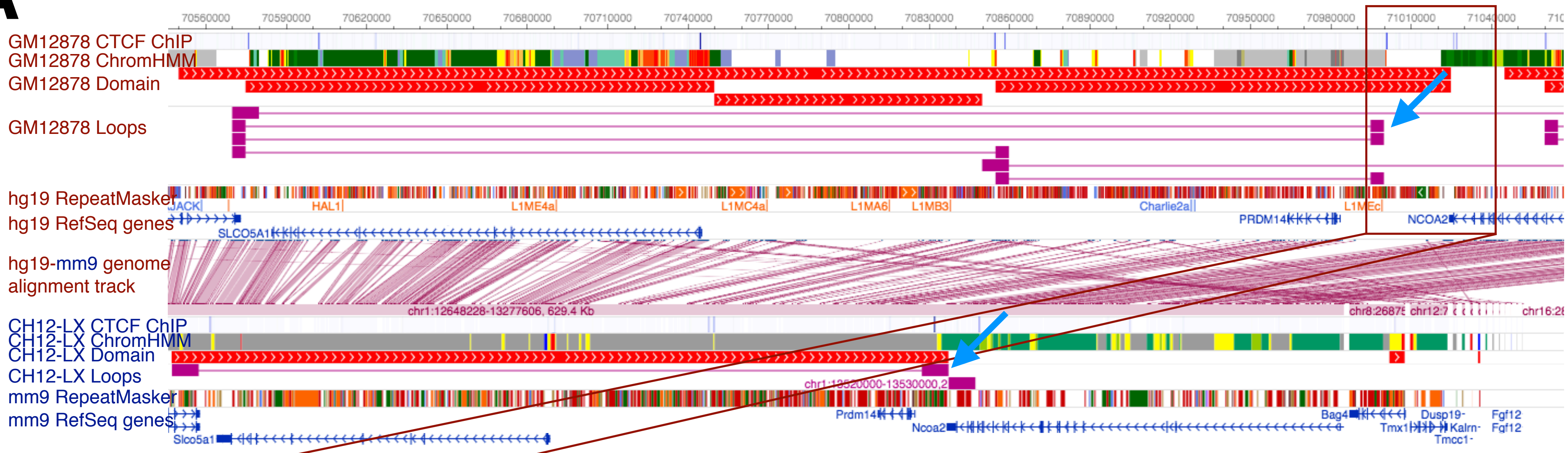


**B**

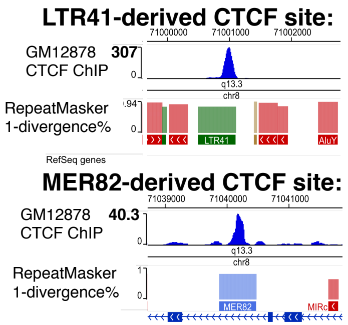


**C**

| Domain Type | Genotype | Percentage of interactions (100kb resolution) Intra-Domain 1 | Intra-Domain 2 | Inter-Domain |
|---|---|---|---|---|
| Targeted Domain | **LTR41-DKO** | **40.1** | **43.1** | **16.9** |
| | WT | 41.5 | 44.5 | 14.1 |
| Negative Control | LTR41-DKO | 47.5 | 39.3 | 13.2 |
| | WT | 48.7 | 38.7 | 12.6 |

1.2-fold
1.1-fold
1.1-fold

**Supplementary Figure 4 | Landscape of the higher-order chromosomal structures conserved in human and mouse maintained by an LTR41-derived CTCF binding site turnover event in humans.** (**A**) Epigenome browser screenshot displaying the conserved genomic and epigenomic landscape. The conserved domain border and loop anchor in human and mouse is marked by blue arrows.(**B**) CTCF ChIP-seq centered on LTR41 and the ancestral MER82. The ancestral MER82 has reduced CTCF binding strength. Note the y-axis upper limits for LTR41 CTCF ChIP-seq track is 307, whereas for MER82 is 40.3 (7.6 times lower). (**C**) Percentage of cross-domain long-range interactions across the targeted domain and a control domain, using Hi-$C^2$ data from LTR41-DKO GM12878 and WT GM12878 cell line

*Visualization from WashU Epigenome Browser: http://epigenomegateway.wustl.edu*

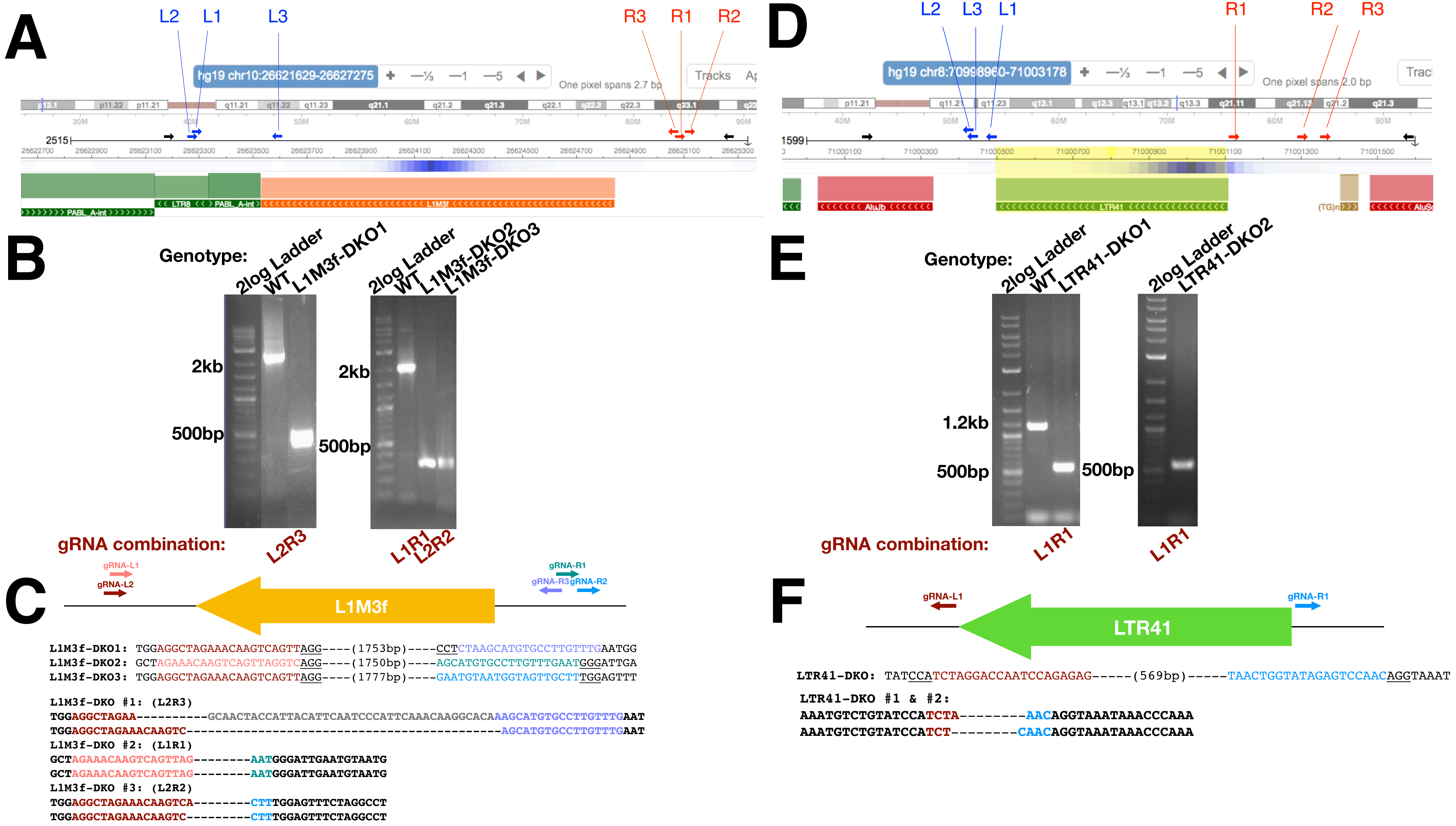


**Supplementary Figure 5 | Experimental design and results of CRISPR-Cas9 mediated deletions of L1M3f and LTR41.** (**A**) Epigenome browser screenshot displaying the exact location of gRNAs pairs (L1-R1, L2-R2, L3-R3) targeting L1M3f and (**D**) LTR41. gRNAs upstream of the TEs are in blue and downstream are in red, cloned into the Cas9-BFP and Cas9-mCherry vectors, respectively). Amplification and Sanger sequencing primers are in black. (**B**) Agarose gel electrophoresis images of PCR amplicons corresponding to the WT and DKO genotypes generated by the CRISPR experiments targeting L1M3f and (**E**) LTR41. (**C**) The CRISPR-mediated genetic breaks are illustrated for L1M3f and (**F**) LTR41 CRISPR clones from Sanger Sequencing traces. gRNA sequences are illustrated with various colors.

*Visualization from WashU Epigenome Browser: http://epigenomegateway.wustl.edu*

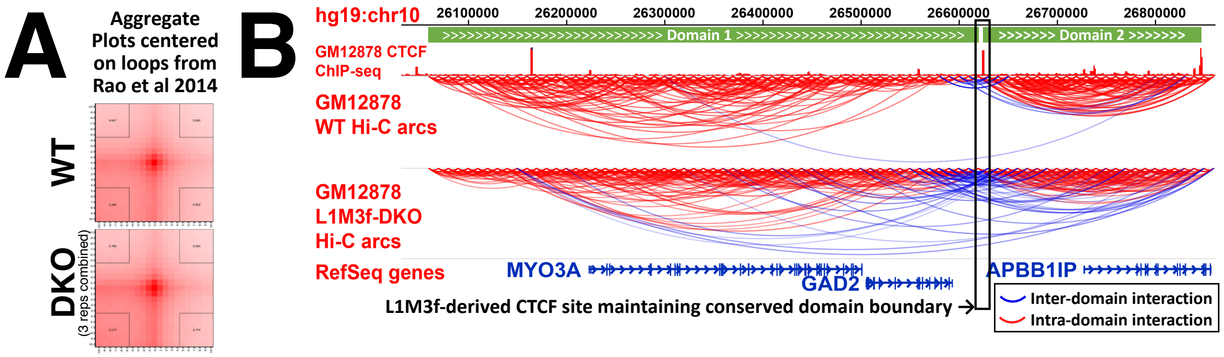


C

| Domain Type | Genotype | Percentage of interactions: Intra-Domain 1 | Intra-Domain 2 | Inter-Domain | | Percentage of interactions (>=25kb): Intra-Domain 1 | Intra-Domain 2 | Inter-Domain | |
|---|---|---|---|---|---|---|---|---|---|
| Targeted Domain | L1M3f-DKO | 63.62 | 23.20 | 13.18 | 2.1x | 56.35 | 16.90 | 26.75 | 1.6x |
| | WT | 65.70 | 27.99 | 6.31 | | 59.58 | 23.96 | 16.46 | |
| Negative Control | L1M3f-DKO | 72.92 | 19.93 | 7.16 | | 70.66 | 10.37 | 18.97 | |
| | WT | 72.09 | 20.38 | 7.53 | | 72.09 | 10.34 | 18.70 | |

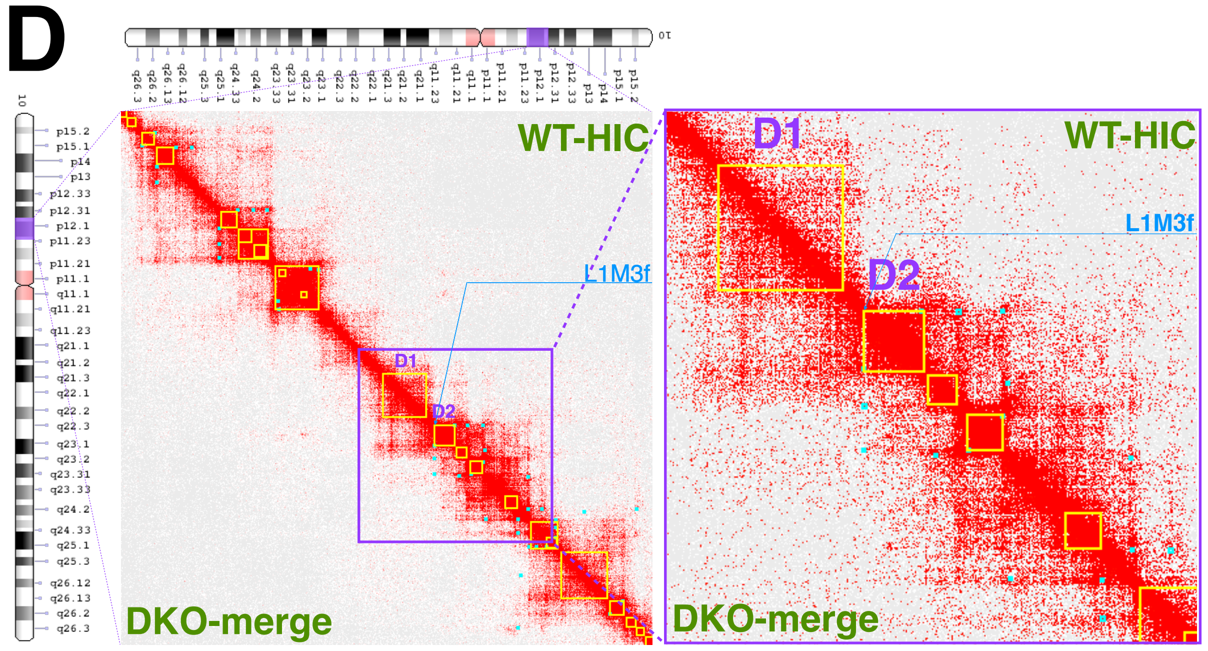


**Supplementary Figure 6 | L1M3f-derived CTCF site is required for maintaining conserved boundary and fidelity of long-range interactions.** (**A**) Aggregate Plot Analysis using two HiC datasets: (i) wild type (WT): highest resolution GM12878 contact map downsampled to 750M reads, and (ii) L1M3f-double knockout (DKO): 3 independent L1M3f-DKO clones were used to generate one HiC library each that was sequenced from 172-333M reads, for a total of ~750M reads. (**B**) Arc view of WT and L1M3f-DKO HiC maps (~750M reads each) shows increased inter-domain interactions (blue arcs) across the domain border that was maintained by the L1M3f element in humans. (**C**) Percentage of intra-domain and inter-domain interactions (left) in WT and L1M3f-DKO GM12878 cells across the targeted domain and a control domain. (right) Percentage of intra-domain and inter-domain long-range interactions (>=25kb) in WT and L1M3f-DKO GM12878 cells across the targeted domain and a control domain. (**D**) Contact map showing the same data as (**B**), but as a heatmap with top-right displaying HiC data generated from WT GM12878 cells and bottom-left displaying HiC data generated from L1M3f-DKO GM12878 cells. Yellow squares and blue dots are domains and loop calls from Rao et al, 2014.

*Visualization from WashU Epigenome Browser: http://epigenomegateway.wustl.edu and Juicebox*

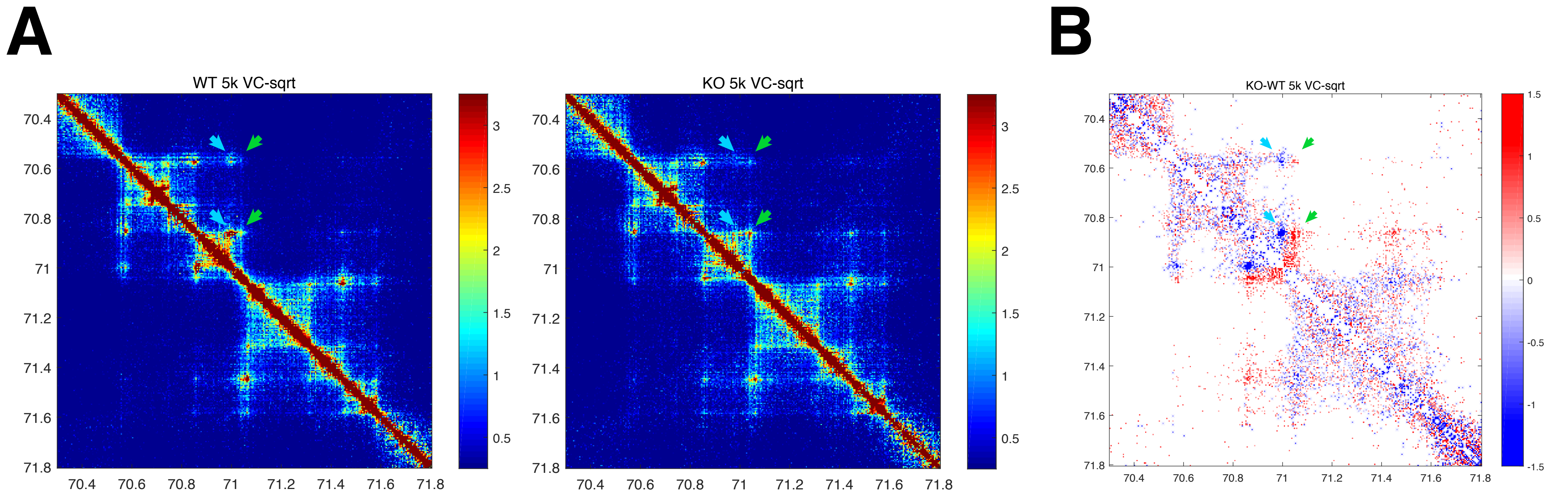


**Supplementary Figure 7 | CTCF binding site turnover from an LTR41-derived CTCF motif to MER82-derived CTCF site.** (**A**) Mega-contact maps generated in Hi-C² experiments for the WT (left) and LTR41-KO (right) locus over a 1.5Mb region (chr8:70.3–71.8 Mb) in GM187278 cells. Blue and green arrows represent the loop's original right anchor (mainly used in WT) and the turned-over loop anchor (mainly used in KO), respectively. (**B**) Differential contact map calculating the difference between the VC-sqrt normalized observed interaction scores of the KO and the WT contact maps. Differential normalized interaction scores for which fold change (KO/WT) > 1.45 or (WT/KO) > 1.45 (fold cut-off), and normalized interaction score of >0.75 (score cut-off) in KO and WT were plotted. Focal blue enrichment suggests loss of loop at the LTR41-derived CTCF loop anchor in the KO as compared to the WT (highlighted by blue arrows). Similarly, focal red enrichment suggests gain of loop at the MER82-derived CTCF loop anchor in KO as compared to the WT (highlighted by green arrows).

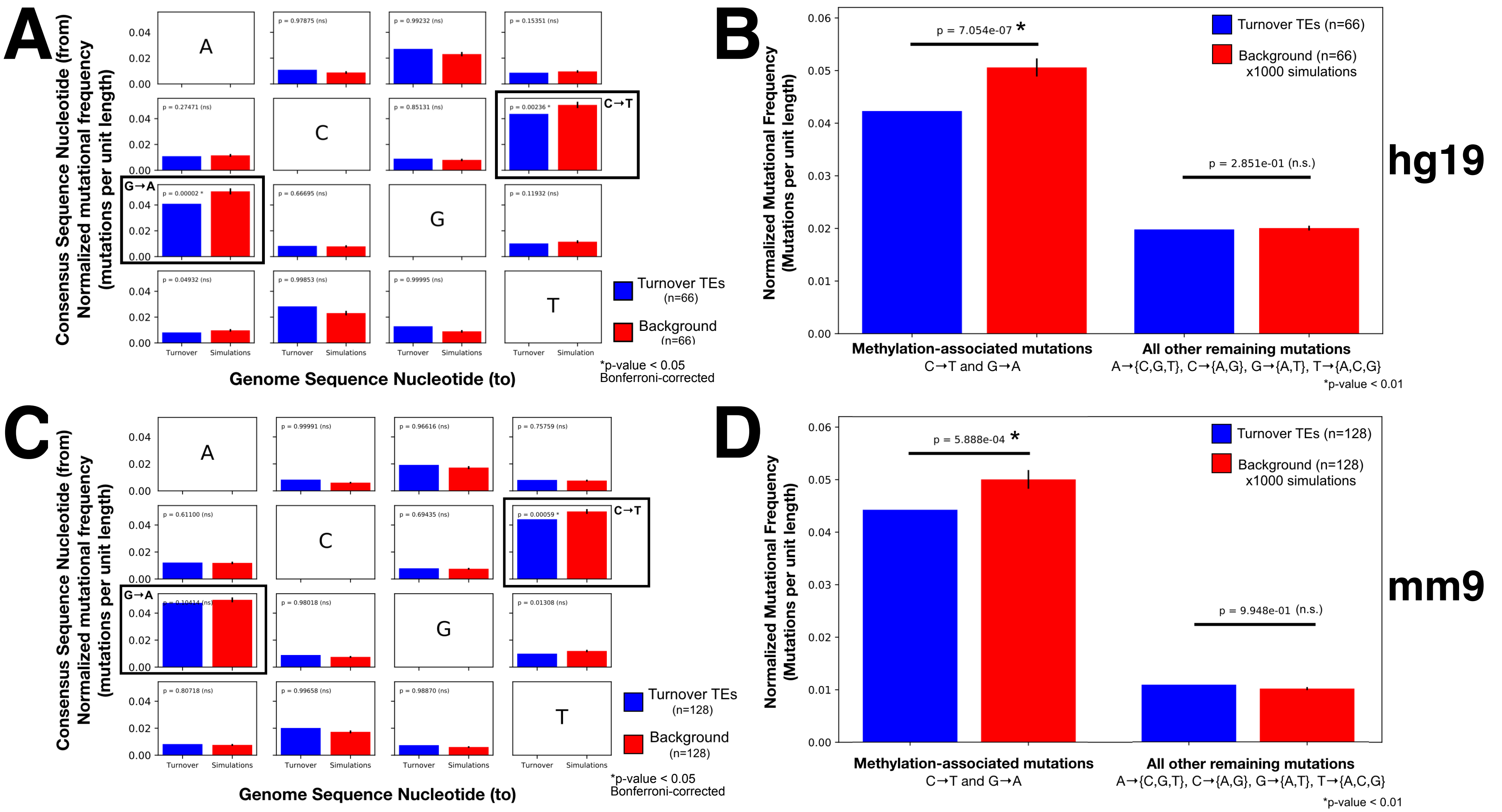


**Supplementary Figure 8 | TEs that potentiate loop anchor turnover in human and mouse show mutational signatures of hypomethylation through evolutionary time (related to Fig 4B):** (**A**) Normalized (by genomic length) mutational frequency of TEs relative to the ancestral human TE sequence using crossmatch alignments. The mean of normalized mutational frequency for TEs involved in turnover (in blue) and the background distribution by repeating the analysis (described in Supplemental Methods) on 1000 permutations of all genomic TEs not involved in looping (in red). Error bars are one standard deviation of the mean from 1000 simulations. (**B**) Methylation-associated mutation frequency in humans computed by taking the average of the C-to-T and G-to-A rates for each TE and averaging over turnover events (in blue) and background (in red). The non-methylation substitution rate was computed by taking the average of all other (ten) single nucleotide substitutions for each TE and then averaging over turnover events (in blue) and background (in red). (**C**) Normalized (by genomic length) mutational frequency of TEs relative to the ancestral mouse TE sequence using crossmatch alignments. (**D**) Methylation-associated mutations and non-methylation-associated mutations in mouse.

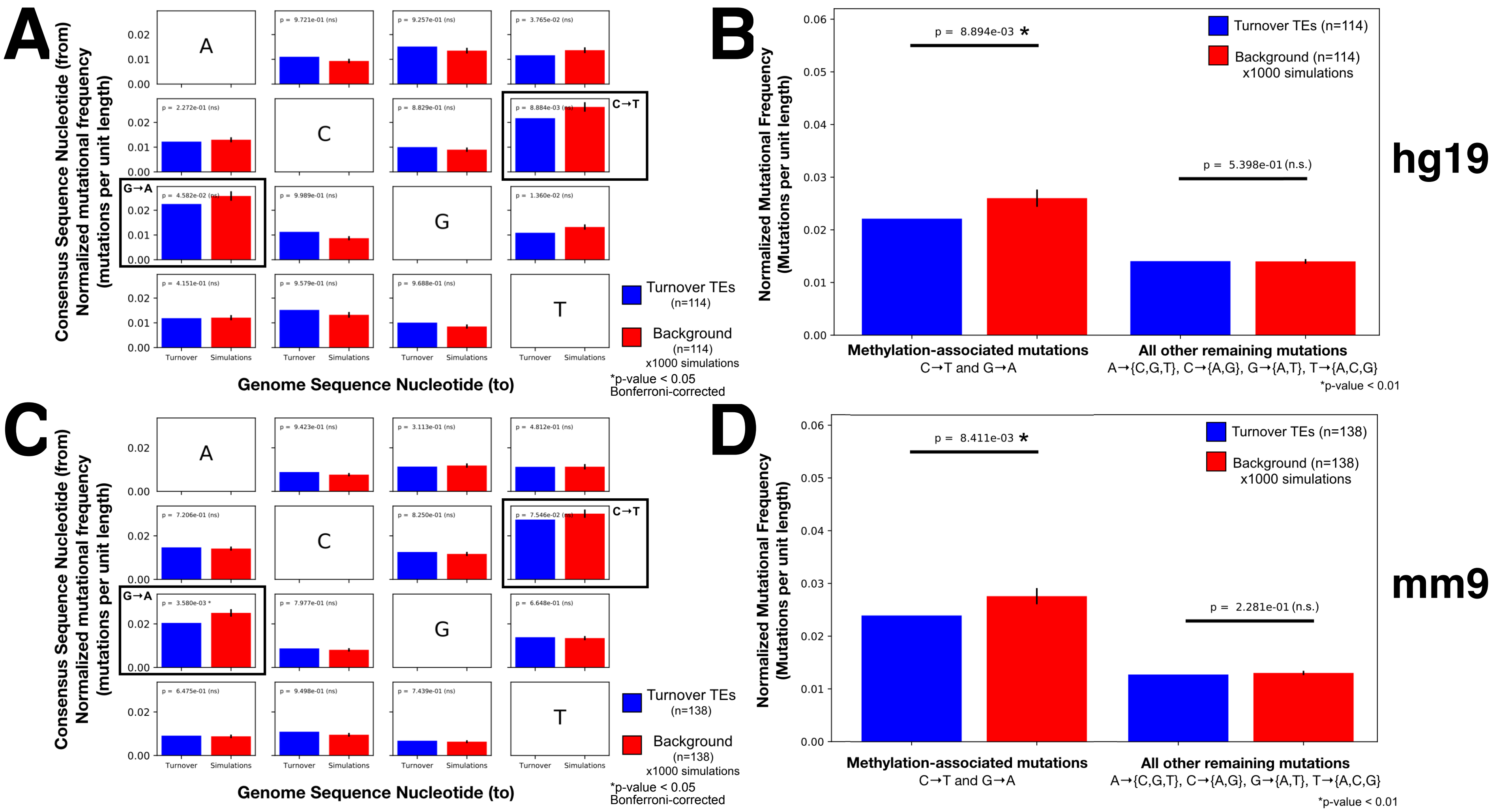


**Supplementary Figure 9 | TEs that potentiate loop anchor turnover in human and mouse show mutational signatures of hypomethylation through evolutionary time (related to Fig 4B):** (**A**) Normalized (by genomic length) mutational frequency of TEs relative to the ancestral human TE sequence using Needle realignments. The mean of normalized mutational frequency for TEs involved in turnover (in blue) and the background distribution by repeating the analysis (described in Supplemental Methods) on 1000 permutations of all genomic TEs not involved in looping (in red). Error bars are one standard deviation of the mean from 1000 simulations. (**B**) Methylation-associated mutation frequency in humans computed by taking the average of the C-to-T and G-to-A rates for each TE and averaging over turnover events (in blue) and background (in red). The non-methylation substitution rate was computed by taking the average of all other (ten) single nucleotide substitutions for each TE and then averaging over turnover events (in blue) and background (in red). (**C**) Normalized (by genomic length) mutational frequency of TEs relative to the ancestral mouse TE sequence using Needle realignments. (**D**) Methylation-associated mutations and non-methylation-associated mutations in mouse.
